# Transcriptomic analysis of resistant and susceptible responses in a new model root-knot nematode infection system using *Solanum torvum* and *Meloidogyne arenaria*

**DOI:** 10.1101/2021.04.02.438176

**Authors:** Kazuki Sato, Taketo Uehara, Julia Holbein, Yuko Sasaki-Sekimoto, Pamela Gan, Takahiro Bino, Katsushi Yamaguchi, Yasunori Ichihashi, Noriko Maki, Shuji Shigenobu, Hiroyuki Ohta, Rochus B. Franke, Shahid Siddique, Florian M. W. Grundler, Takamasa Suzuki, Yasuhiro Kadota, Ken Shirasu

**Author notes:** Corresponding authors: Yasuhiro Kadota, Ken Shirasu.

## Abstract

Root-knot nematodes (RKNs) are among the most devastating pests in agriculture. *Solanum torvum* Sw. (turkey berry) has been used as a rootstock for eggplant (aubergine) cultivation because of its resistance to RKNs, including *Meloidogyne incognita* and *M. arenaria*. We previously found that a pathotype of *M. arenaria*, A2-J, is able to infect and propagate in *S. torvum. In vitro* infection assays showed that *S. torvum* induces the accumulation of brown pigments during avirulent pathotype A2-O infection, but not during virulent A2-J infection. This experimental system is advantageous because resistant and susceptible responses can be distinguished within a few days, and because a single plant genome can yield information about both resistant and susceptible responses. Comparative RNA-sequencing analysis of *S. torvum* inoculated with A2-J and A2-O at early stages of infection was used to parse the specific resistance and susceptible responses. Infection with A2-J did not induce statistically significant changes in gene expression within one day post-inoculation (DPI), but afterward, A2-J specifically induced the expression of chalcone synthase, spermidine synthase, and genes related to cell wall modification and transmembrane transport. Infection with A2-O rapidly induced the expression of genes encoding class III peroxidases, sesquiterpene synthases, and fatty acid desaturases at 1 DPI, followed by genes involved in defense, hormone signaling, and the biosynthesis of lignin at 3 DPI. Both isolates induced the expression of suberin biosynthetic genes, which may be triggered by wounding during nematode infection. Histochemical analysis revealed that A2-O, but not A2-J, induced lignin accumulation at the root tip, suggesting that physical reinforcement of cell walls with lignin is an important defense response against nematodes. The *S. torvum*-RKN system can provide a molecular basis for understanding plant-nematode interactions.

## INTRODUCTION

Plant-parasitic nematodes (PPNs) infect a broad range of commercially important crop families such as the Solanaceae (tomato, potato, pepper), Fabaceae (soybean, lucerne, lentils), Malvaceae (cotton), Amaranthaceae (sugar beet), and Poaceae (syn. Gramineae; rice, wheat, maize), causing an estimated annual loss of $80 billion USD (Jones et al., 2013; Sato et al., 2019). The most economically important group of PPNs are sedentary endoparasites, including root-knot nematodes (RKNs) and cyst nematodes (CNs) (Palomares-Rius et al., 2017). Sedentary endoparasites induce the formation of permanent feeding cells that provide specialized nutrient sources for nematodes (Bartlem et al., 2014; Siddique and Grundler, 2018; Smant et al., 2018). Infective second-stage juveniles (J2s) of RKNs (*Meloidogyne* spp.) predominantly invade near the root tip and then migrate intercellularly toward the apical meristematic region without crossing the endodermis, making a U-turn to enter the vascular cylinder where they induce several giant cells as a feeding site by stimulating the redifferentiation of root cells into multinucleate giant cells by repeated nuclear divisions without cytoplasmic division. After maturation, adult RKN females lay eggs in a gelatinous egg mass on or below the surface of the root (Sijmons et al., 1991; Abad et al., 2009; Escobar et al., 2015). In contrast, CNs move destructively through cells into the vascular cylinder, select a single cell, and form a syncytium as a feeding site by local dissolution of cell walls and protoplast fusion of neighboring cells. A CN female produces hundreds of eggs and its body forms a cyst that can protect the eggs for many years in the soil (Wyss and Zunke, 1986; Bohlmann and Sobczak, 2014; Bohlmann, 2015).

Nematicides have been commonly used to control PPNs in agriculture, but some nematicides such as methyl bromide and aldicarb are currently banned from use in many countries due to their negative effects on the environment and human health (Zasada et al., 2010; Brennan et al., 2020; Oka, 2020). It has therefore become important to understand the molecular mechanisms of plant immunity against PPNs to provide a foundation for the development of new environmentally-friendly and effective control methods.

In general, the plant immune system is represented by two inter-related tiers (Jones and Dangl, 2006; Dodds and Rathjen, 2010). The first is governed by cell surface-localized pattern recognition receptors (PRRs) that perceive pathogen-associated molecular patterns (PAMPs), leading to pattern-triggered immunity (PTI) (Boutrot and Zipfel, 2017). Successful pathogens secrete effector molecules into the apoplast or directly into plant cells, which interfere with PTI, resulting in successful infection. Resistant plants recognize cell-invading effectors through recognition by intracellular nucleotide-binding domain leucine-rich repeat (NLR)-type immune receptors, which are encoded by resistance (*R*) genes. Similar mechanisms are also conserved in plant-PPN interactions. For example, the well-conserved nematode pheromone ascaroside has been identified as a PAMP (Manosalva et al., 2015), but the corresponding PRR has not yet been found. PPN genome sequence analyses identified a number of candidate virulence effectors (summarized in Mejias et al. (2019)), and a handful of NLR protein-encoding *R* genes involved in PPN recognition have been well-studied and characterized, including tomato *Mi-1.2, Mi-9*, and *Hero-A*; potato *Gpa2* and *Gro1-4*; pepper *CaMi*; and prune *Ma* (Milligan et al., 1998; van der Vossen et al., 2000; Ernst et al., 2002; Paal et al., 2004; Chen et al., 2007; Jablonska et al., 2007; Claverie et al., 2011; Kaloshian et al., 2011). *Mi-1.2, Mi-9, CaMi*, and *Ma* confer resistance against RKNs, whereas *Hero-A, Gpa2*, and *Gro1-4* provide resistance against CNs.

Although the PPN perception mechanism is somewhat clearer at the molecular level, it is still largely unknown what kind of downstream responses are induced after the recognition of avirulent PPNs. It is also unclear what kind of host responses are induced after infection with virulent PPNs, leading to susceptibility and infestation. There are several difficulties in working on plant responses against PPNs. First and foremost, most model plants, such as Arabidopsis, are susceptible to PPNs and therefore cannot be used to study the cascade of responses leading to resistance. Second, PPNs migrate long-distances inside roots, inducing complicated responses as they go, triggered by mechanical stress and wounding, among others, making it difficult to pinpoint the key genes involved in resistance or susceptibility by transcriptome analyses. Some studies have used comparative transcriptomics using susceptible and resistant plants infected with a single genotype of nematode (Postnikova et al., 2015; Xing et al., 2017; Ye et al., 2017; Zhang et al., 2017a). However, it is difficult to rule out the possibility that differences in gene expression were due to resistance or susceptibility rather than to differences in the genetic backgrounds of host plants. Lastly, susceptible responses such as the formation of feeding sites are induced in specific cells targeted by PPNs, and defense responses are likely to be induced in the cells directly impacted by PPN activity. Thus, cells responding to PPNs are rather limited, making the analysis technically challenging.

Here we have introduced *Solanum torvum* Sw “Torvum Vigor” to overcome these problems. *S. torvum* has been widely used as a rootstock of eggplant (aubergine) to prevent disease caused by PPNs, as well as the soil-borne pathogens *Ralstonia solanacearum, Verticillium dahliae*, and *Fusarium oxysporum* f. *melongenae* n. f. (Gousset et al., 2005; Yamaguchi et al., 2010; Bagnaresi et al., 2013; Yang et al., 2014; Uehara et al., 2017; García-Mendívil et al., 2019; García-Mendívil and Sorribas, 2019; Murata and Uesugi, 2021). *S. torvum* Sw “Torvum Vigor” is resistant to *Meloidogyne arenaria* pathotype A2-O (A2-O), but susceptible to *M. arenaria* pathotype A2-J (A2-J) (Uehara et al., 2017). By using *S. torvum* and avirulent or virulent isolates, we established an *in vitro* infection system and performed comparative transcriptome analyses to identify genes whose expressions were associated with either resistance or susceptibility by carefully collecting only root tips attacked by RKNs, which allowed us to detect gene expression only in cells directly affected by nematodes. In addition, observation of infected root tip morphology suggests that the success or failure of the immune system against PPNs is determined within a few days of invasion. Thus, we decided to focus on the transcriptional changes that occurred in the very early stages of infection, which has not been studied in previous transcriptomic analyses (Bagnaresi et al., 2013; Postnikova et al., 2015; Xing et al., 2017).

Comparative clustering analyses of differentially expressed genes (DEGs) identified a large number of novel genes, especially those involved in susceptibility through cell wall modification and transmembrane transport; resistance through lignin and isoprenoid biosynthesis and fatty acid metabolism; and suberin biosynthesis in mechanical wounding. Consistent with the transcriptional up-regulation of lignin biosynthetic genes from A2-O invasion, lignin is accumulated at the root tip of *S. torvum* infected with avirulent A2-O but not with virulent A2-J, suggesting that *S. torvum* reinforces the cell wall as a defense response against the avirulent RKN.

## MATERIALS AND METHODS

### Plant Materials and Growth Conditions

Seeds of *S. torvum* cultivar “Torvum Vigor” were sown on half-strength Murashige-Skoog (MS) medium containing 1% sucrose. Plants were grown in a controlled growth chamber under long-day photoperiods (16 hours light/8 hours dark) at 25 °C.

### Nematode Infection Assay

*M. arenaria* pathotypes A2-J and A2-O were propagated on *Solanum lycopersicum* cultivar “Micro-Tom” in a greenhouse. Nematode eggs were isolated from infected roots and then hatched at 25 °C. Freshly hatched J2s were collected and transferred to a Kimwipe filter (a folded Kimwipe tissue) placed on the top of a glass beaker filled with sterilized distilled water (SDW) containing 100 μg/ml streptomycin and 10 μg/ml nystatin. Only active J2s pass through the filter. Filtered J2s were surface sterilized with 0.002 % mercuric chloride, 0.002 % sodium azide, and 0.001 % Triton X-100 for 10 min, and then rinsed three times with SDW (Mitchum et al., 2004). Eleven-day-old *S. torvum* seedlings grown in 6-well plates were inoculated with 200-300 J2s resuspended in SDW. The plates were wrapped in aluminum foil for 2-3 days after inoculation to promote nematode infection. When mature giant cells were observed 18 days post-inoculation (DPI), we used the MS media without sucrose to prevent the formation of callus-like structures. The difference in the number of normal galls formed by A2-J or A2-O at 4 DPI was statistically tested using the Mann-Whitney *U* test with R software (v3.6.3).

Nematodes resident in root tissues were stained with acid fuchsin 2-4 DPI (Bybd et al., 1983), photographed by light microscopy (Olympus BX51), and the photomicrographs were processed using cellSens (Olympus, Japan). For the observation of giant cells and developing nematodes at 18 DPI, infection sites were fixed with glutaraldehyde and cleaned with benzyl-alcohol/benzyl-benzoate (BABB) (Cabrera et al., 2018). We observed BABB-cleaned samples by confocal laser scanning microscopy (TCS SP5, Leica Microsystems GmbH, Germany). Photomicrographs were processed using LAS X software (version 3.7.0.20979, Leica Microsystems GmbH).

### RNA-sequencing (RNA-seq) and *De Novo* Assembly

*S. torvum* seedlings were grown on half-strength MS medium containing 1% sucrose. Eleven-day-old seedlings were treated with SDW as a mock infection or with 200-300 J2s of *M. arenaria* A2-J for susceptible infection or A2-O for resistant infection. Root tips attacked by the nematodes were checked under microscopy, and more than 50 root tips were cut (approximately 3-5 mm from the tip) and collected for each treatment (Figure 2A). Root tip samples were collected at 1, 2, and 3 DPI with four biological replicates. Whole shoot and root samples were collected at 1, 3, 6, and 9 DPI with four biological replicates. Root tip samples were used for *de novo* assembly and differential gene expression analyses, and whole shoot and root samples were used only for *de novo* assembly.

RNA-seq libraries were prepared from the collected samples using a high-throughput RNA-seq method (Kumar et al., 2012). The 85-bp paired-end reads for the root tip samples, and the 85-bp single-end reads for the whole shoot and root samples were sequenced on an Illumina NextSeq 500 platform (Illumina, CA, USA). The FASTX toolkit 0.0.13.2 (Hannonlab) was used for quality filtering. Low-quality nucleotides (Phred scores of <30) were removed from the 3’ ends, and short reads (<76bp) were excluded. Reads with at least 95% of nucleotides with Phred scores>20 were kept and used for the downstream analyses (Supplementary Table 1A). Adaptor sequences were removed using custom scripts written in Perl (Kumar et al., 2012). Filtered reads were mapped to the genome assembly of *M. arenaria* A2-J (GenBank accession number JAEEAS010000000) or A2-O (GenBank accession number QEUI01000000) (Sato et al., 2018) using HISAT2 (version 2.1.0) (Kim et al., 2019) to exclude reads of nematode origin. Unmapped reads were used for *de novo* transcriptome assembly (Supplementary Figure 1A).

Three different transcriptome assemblers were used for *de novo* assembly: SOAPdenovo-Trans v1.03 (Xie et al., 2014), Velvet v1.2.10 (Zerbino and Birney, 2008)/Oases v0.2.09 (Schulz et al., 2012) and Trinity package v2.4.0 (Grabherr et al., 2011; Haas et al., 2013). Unmapped paired-end and single-end reads were normalized using Trinity and assembled independently (Mamrot et al., 2017). For assemblies of each dataset, SOAPdenovo-Trans was set at *k*-mer sizes: 21, 23, 25, 27, 29, 31, 35, 41, 51, 61, Velvet/Oases was set at *k*-mer sizes: 21, 23, 25, 27, 29, 31, 35, 41, 51, 61, and Trinity was set at *k*-mer size 25 (Supplementary Table 1B).

Assemblies were merged into a non-redundant dataset using the EvidentialGene pipeline (http://arthropods.eugenes.org/genes2/about/EvidentialGenetrassemblypipe.html,version 2017.12.21) as previously described (Nakasugi et al., 2014). Oases assembled scaffolds were split at gaps into contigs before merging with contigs from the other assemblies with the EvidentialGene tr2aacds pipeline. The tr2aacds pipeline produces ‘primary’ and ‘alternate’ sequences of non-redundant transcripts with ‘primary’ transcripts being the longest coding sequence for a predicted locus. Next, we used the evgmrna2tsa program from EvidentialGene to generate mRNA, coding, and protein sequences. BUSCO v3.0.2 (Simão et al., 2015) was applied for quantitative assessment of assembly completeness. This assembly and one previously reported for *S. torvum* by Yang et al. (2014) (GenBank accession number GBEG01000000) were compared to the Embryophyta odb9 dataset, which contains 1,440 BUSCO groups. The homology of the contigs from the final assembly was searched against the NCBI non-redundant database using BLASTX (BLAST+ v2.7.1) with an e-value threshold of 1E-05. We also compared the contigs with Arabidopsis genome annotation (TAIR10) (Berardini et al., 2015) using BLASTX at the e-value cutoff of 10. Results of the annotation are summarized in Supplementary Table 2.

### Differential Expression Analysis

Quantification of *S. torvum* transcripts was performed with the mapping-based mode of Salmon v0.10.2 (Patro et al., 2017) by using reads that did not map to nematode genome assemblies. Quantified transcript-level abundance data were imported to R (v3.6.3) using tximport v.1.14.2 (Soneson et al., 2015) package, and differential gene expression analysis was carried out with the edgeR package (v3.28.1) (Robinson et al., 2010). Transcripts with very low expression values were filtered out with the “filterByExpr” function. Differentially expressed genes (DEG) (false discovery rate (FDR) ≤ 0.01, log-transformed fold change (logFC) ≥ 1 or logFC ≤ −1) were identified using the quasi-likelihood F-test by comparing expression levels during infection with A2-J or A2-O to mock treatment at the same time point. Counts per million genes and DEGs are listed in Supplementary Tables 3 and 4, respectively.

### Principal Component Analysis (PCA) with Self-Organizing Map (SOM) Clustering

To group genes by expression pattern, we applied the SOM clustering method on genes within the top 25 % of the coefficient of variation for expression across samples as previously described (Chitwood et al., 2013; Ranjan et al., 2014; Goto et al., 2020). Scaled expression values, representing the average principal component (PC) values among each gene in a cluster were used for multilevel three-by-three hexagonal SOM (Wehrens and Buydens, 2007). One hundred training iterations were used during the clustering, over which the α learning rate decreased from 0.0071 to 0.0061. The final assignment of genes to winning units formed the basis of the gene clusters. The results of SOM clustering were visualized in a PCA space where PC values were calculated based on gene expression across samples (Figure 4A).

### Functional Annotation of Transcriptome and Gene Ontology (GO) Enrichment Analysis

We compared the contigs of our assembly with the NCBI non-redundant database using BLASTX (BLAST+ v2.7.1) with an e-value threshold of 1E-05. In addition, predicted amino acid sequences that begin with methionine were also annotated using InterProScan (v5.32-71.0) (Jones et al., 2014). BLASTX and InterProScan outputs were used for Blast2GO (v5.2.5) analysis to annotate the contigs with GO terms (Gotz et al., 2008). GO enrichment analyses of the sets of genes induced by A2-O infection at 1 DPI or that were assigned to each cluster generated by SOM was performed by comparison with all genes using GO terms generated by Blast2GO at the FDR cutoff of 1E-04 (Gotz et al., 2008). We further used the “Reduce to most specific terms” option in Blast2GO to remove general GO terms and obtain only the most specific GO terms.

### Histochemistry of Lignin Deposition

Lignin deposition during infection with A2-J or A2-O was visualized by phloroglucinol-HCl staining as previously described (Jensen, 1962). Eleven-day-old *S. torvum* seedlings were inoculated with nematodes or treated with SDW (mock) as described above. We collected root tips 3 DPI for lignin staining with phloroglucinol-HCl. Microphotographs were taken and processed as described above and combined manually.

### Aliphatic Suberin Monomer Analysis

Quantification of aliphatic suberin was performed as described previously (Holbein et al., 2019). Eleven-day-old plants were treated with SDW (mock) or infected with A2-J or A2-O. At 4 DPI, root tips inoculated with nematodes were microscopically checked for infection, and more than 50 infected root tips were cut (approximately 3-5 mm from the tip) and collected for each treatment. To remove unbound lipids, samples were extracted in methanol for 24 h then in chloroform for 24 h, dried, and weighed. Samples were depolymerized and analyzed by GC-MS (Agilent 6890N-Agilent 5973N quadrupole mass-selective detector, Agilent Technologies, Germany) for monomer identification and for quantitative analysis based on an internal standard using an identical GC system coupled with a flame ionization detector as described previously (Franke et al., 2005).

## RESULTS

### A2-O induces a Browning Response, and A2-J induces Gall Formation in *S. torvum*

To understand the differential responses of *S. torvum* to *M. arenaria* A2-J and A2-O, we first established an *in vitro* infection system. Seedlings of *S. torvum* were grown in MS agar plates for 11 days and then inoculated with 200-300 J2s of A2-J or A2-O. At 4 DPI, more than 90 % of root tips infected with A2-J induced the formation of gall-like structures ranging in size. These galls are classified here as “normal” galls, while the rest produced brown pigments. Normal galls lacked obvious brown pigment accumulation and were further classified based on the width of the gall into small (shorter than 0.5 mm), medium (0.5-0.8 mm), and large (wider than 0.8mm) (Figure 1A). In contrast, about 60 % of A2-O-infected root tips accumulated at least some brown pigment. Some of these brownish root tips also had an abnormal appearance due to the formation of balloon-like structures, and others had many localized and highly pigmented spots. There were a very few small gall-like structures formed after infection with A2-O, but far fewer and smaller than in root tips infected with A2-J (Figure 1B). RKN staining by acid fuchsin revealed that both A2-J and A2-O successfully invaded the roots (Figure 1C). Interestingly, host cells invaded by A2-O uniformly accumulated brownish pigments, suggesting that the surrounding tissue is strongly responding to, and highly correlated with A2-O infection, a response that was absent from A2-J infected roots. It is generally known that browning of plant tissue is related to enzymatic or non-enzymatic oxidation of phenolic substances (Mesquita and Queiroz, 2013), but the identity of the brown pigments synthesized upon infection with A2-O is unknown. By 18 DPI, A2-J had induced the formation of mature multinucleate giant cells and developed into fourth stage juveniles (Figure 1D). In contrast, A2-O did not induce the formation of giant cells nor did it develop past second stage juveniles. These results suggest that *S. torvum* rapidly induces defense responses against A2-O, which inhibits the maturation of A2-O and gall formation. In contrast, A2-J inhibits or evades the induction of defense responses, continues development, and induces gall formation.

**FIGURE 1.**
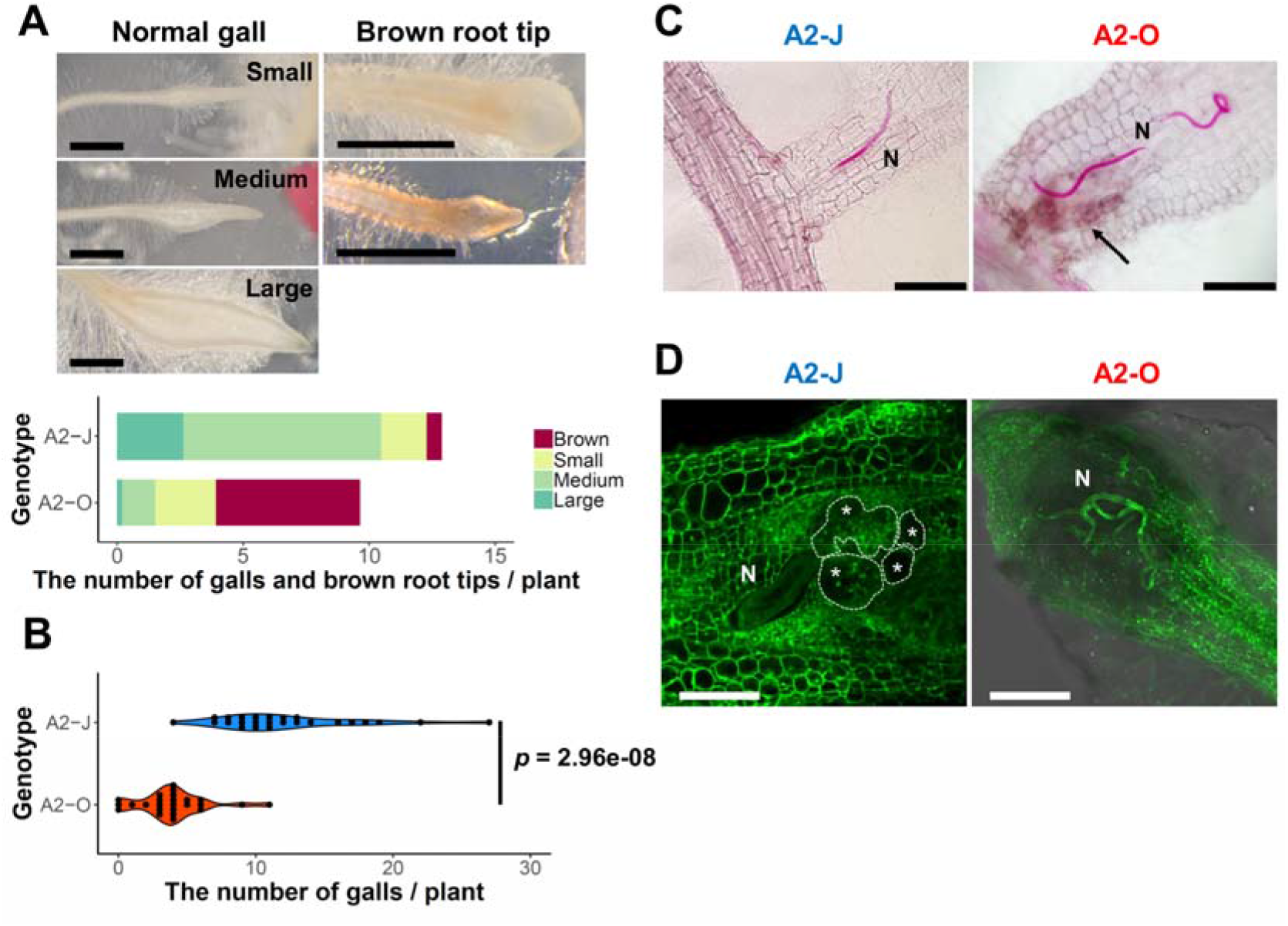
*S. torvum* infection with *M. arenaria* A2-J induced gall formation, and A2-O induced the accumulation of brown pigment at root tips. (A) Microscopic examination of *S. torvum* root tips infected with A2-J or A2-O. Seedlings of *S. torvum* were infected with 200-300 *M. arenaria* A2-J or A2-O juveniles. The number of galls and brown root tips per plant were counted at 4 DPI. Normal galls without accumulation of brown pigment were binned by gall length into small (shorter than 0.5 mm), medium (0.5-0.8 mm), or large (longer than 0.8 mm). Scale bars indicate 1 mm. The average number of galls per plant (*n*= 25) is shown as a bar chart. Experiments were performed four times with similar results. (**B**) A2-J induced the formation of more galls than A2-O. The violin plot indicates variation in the number of galls per plant at 4 DPI. A *p*-value was calculated by the Mann-Whitney *U* test from the mean number of galls per plant. Similar results were obtained from four independent experiments. (**C**) Both A2-J and A2-O entered roots. Nematodes (N) in the roots were stained with acid fuchsin 2-4 DPI. The arrow indicates the accumulation of brown pigment. Scale bars indicate 200 μm. (**D**) *S. torvum* inhibited the growth of A2-O, but not A2-J. Nematodes and giant cells in the *S. torvum* roots 18 DPI were visualized by a method described previously (Cabrera et al., Int. J. Mol. Sci. 2018, 19, 429). Nematodes and giant cells are indicated by N and asterisks, respectively. Scale bars indicate 200 μm.

### RNA-seq Analysis of *S. torvum* Root Tips Infected with A2-O or A2-J

RNA-seq analysis was performed to understand the differences in transcriptional regulation of the *S. torvum* response to infection by nematodes that induce an immune response or that are successful parasites. Eleven-day-old *S. torvum* seedlings grown on MS agar plates were inoculated with 200-300 surface-sterilized J2s of A2-J or A2-O, or treated with SDW (mock treatment) *in vitro*. Since there were clear morphological differences between the root tips infected with A2-J and A2-O after four days (Figure 1A), it should follow that the success or failure of infection is determined within a few days post-inoculation. We therefore decided to analyze the transcriptome at 1-3 DPI, corresponding to the early stages of infection. In addition, to detect gene expression in cells directly affected by the nematodes, we carefully collected infected root tips under a stereomicroscope (Figure 2A). Root tips were cut with precision forceps and flash-frozen with liquid nitrogen to preclude the induction of wound responses. More than 50 root tips were pooled for each treatment, and four biological replicates were used for the RNA-seq based transcriptome analyses. We also carried out RNA-seq of whole roots and shoots of *S. torvum* infected with A2-J or A2-O, or mock treatment (1, 3, 6, and 9 DPI) to improve the completeness of *de novo* transcriptome assembly. As a result, we obtained 218,024,788 paired-end reads from root tips and 341,297,551 single-end reads from whole shoots and roots after quality filtering (Supplementary Table 1A). After removing the reads derived from nematodes, we performed *de novo* assembly using multiple assemblers with a variety of *k*-mer sizes (Supplementary Figure 1A and Supplementary Table 1B). A set of non-redundant transcripts was generated by merging these multiple assemblies. The final assembly had 88,596 contigs with an N50 of 1,298 bp, an average size of 800.62 bp, and a total length of 70,931,593 bp (Table 1). We assessed the accuracy and completeness of the final assembly using BUSCO. The assembly included an estimated > 95 % of the assessed dataset, improving the current status of the transcriptome assembly of *S. torvum* (Yang et al. (2014) (Supplementary Figure 1B) and provided a high-quality transcriptome assembly of *S. torvum* for further analyses.

**FIGURE 2.**
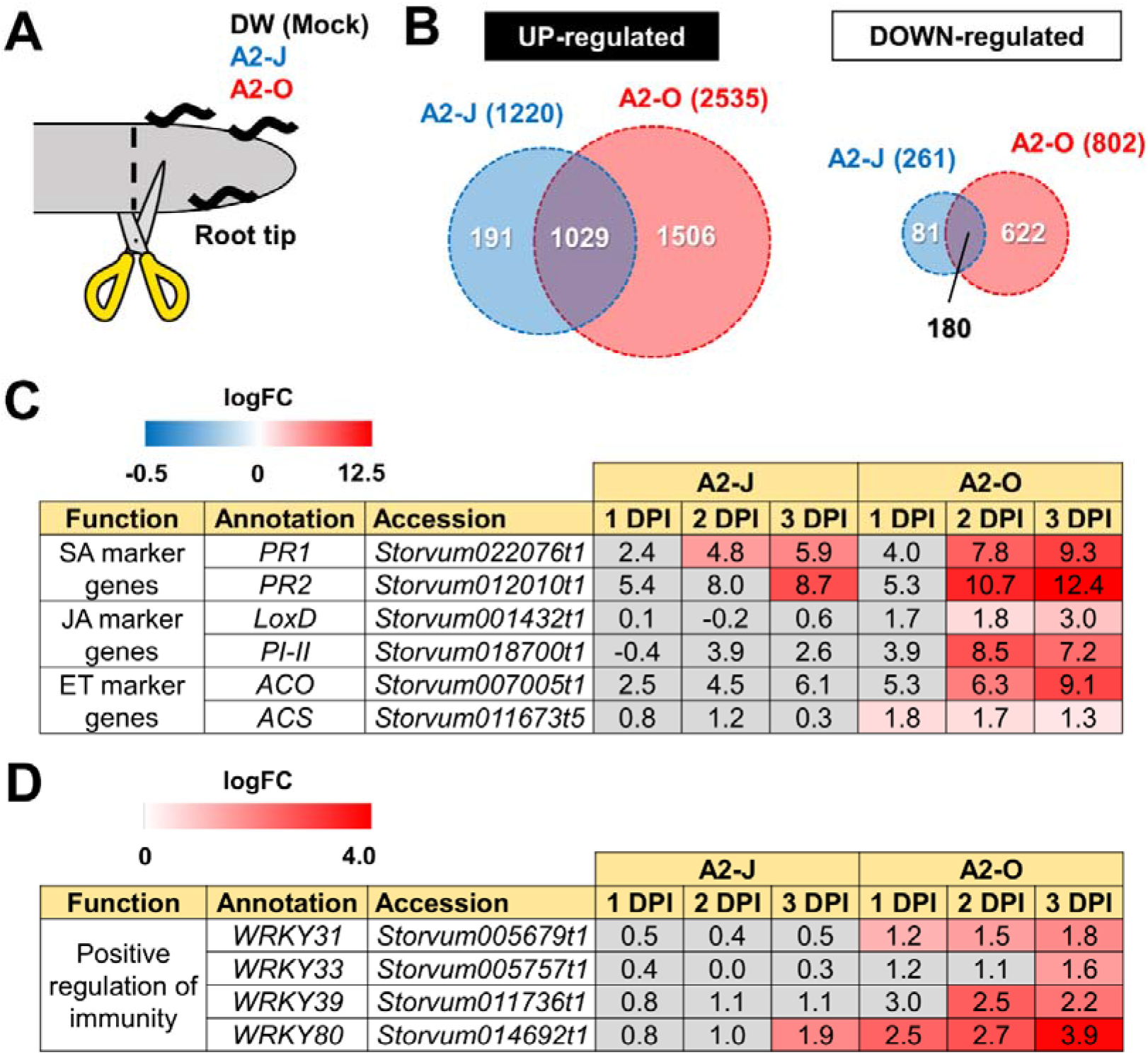
Transcriptome analysis of differentially expressed genes (DEGs) in *S. torvum* infected with *M. arenaria* A2-J or A2-O. (**A**) Sampling of *S. torvum* root tips infected by A2-J or A2-O for RNA-seq analysis. Root tips (approximately 3-5 mm from the tip) were cut (scissors not to scale) and collected at 1, 2, and 3 DPI for RNA-seq. (**B**) Venn diagrams show the overlap of up-regulated genes (logFC ≥ 1, FDR ≤ 0.01) and down-regulated genes (logFC ≤ −1, FDR ≤ 0.01) for at least one time point after infection with A2-J or A2-O. (**C, D**) Infection with A2-O induced the expression of phytohormone marker genes and WRKY transcription factors. *PR1* and *PR2* are marker genes of salicylic acid (SA), *LoxD* and *PI-II* are marker genes for jasmonic acid (JA), and *ACS* and *ACO* are marker genes for ethylene (ET). The homologs of tomato WRKY genes *WRKY31, WRKY33, WRKY39*, and *WRKY80* positively regulate immunity. The logFC values of the genes (compared to mock treatment) at 1, 2, and 3 DPI are shown. Heatmaps indicate relative gene expression levels. Grey color indicates values with no statistically significant differences at |logFC| ≥ 1 and FDR ≤ 0.01.

**TABLE 1.** *De novo* assembly of the *S. torvum* transcriptome

| Parameter |  |
| --- | --- |
| Number of contigs | 88,596 |
| Total size of contigs | 70,931,593 bp |
| Average length | 800.62 bp |
| Median length | 469.00 bp |
| N50 contig length | 1,298 bp |
| Minimum length | 96 bp |
| Maximum length | 16,903 bp |

### Infection with A2-O, but not A2-J, Induces the Expression of Defense-related Genes

Differential expression analysis showed that 1,220 genes were significantly up-regulated and 261 genes were down-regulated upon infection with A2-J, while 2,535 genes were up-regulated and 802 genes were down-regulated by infection with A2-O at at least one-time point during root tip infection, compared to the mock treatment (logFC ≥ 1 for up-regulation and logFC ≤ −1 for down-regulation, FDR ≤ 0.01, Figure 2B; Supplementary Figure 2; Supplementary Table 4). 1,029 genes were up-regulated, and 180 genes were down-regulated at at least one-time point in both A2-J and A2-O infected plants (Figure 2B). Previous studies showed that the expression of genes associated with the salicylic acid (SA), jasmonic acid (JA), and ethylene (ET) signaling pathways are induced in resistant plants infected with PPNs (Ye et al., 2017; Zhang et al., 2017a; Bali et al., 2019; Du et al., 2020; Ghaemi et al., 2020), so we investigated the expression of marker genes for hormone biosynthesis, hormone signaling (Figure 2C), and defense responses (Figure 2D). Infection with A2-O significantly induced the expression of SA-regulated genes that encode basic pathogenesis-related protein 1 (*PR1*) and β-1,3-glucanase (*PR2*), as well as a JA biosynthesis gene (lipoxygenase D (*LoxD*)), the JA-regulated gene protease inhibitor II (*PI-II*), and two ET biosynthesis genes (1-aminocyclopropane-1-carboxylate synthase *(ACS)* and 1-aminocyclopropane-1-carboxylate oxidase *(ACO))* (Farmer et al., 1992; Yan et al., 2013; Booker and DeLong, 2015; Zhang et al., 2018; Marhavý et al., 2019) (Figure 2C). Infection with A2-J induced the expression of *PR1* and *PR2* to a lesser extent, but did not induce the expression of *LoxD, PI-II, ACS*, or *ACO*.

Tomato (*S. lycopersicum*) WRKY transcription factors are induced by pathogen infection and positively regulate defense responses against pathogens (Liu et al., 2014; Li et al., 2015a; b; Zhou et al., 2015). For example, SlWRKY31 and SlWRKY33 play important roles in resistance against fungal and oomycete pathogens, while SlWRKY39 is involved in resistance against bacterial pathogens (Huang et al., 2012; Sun et al., 2015). SlWRKY80 is required for *Mi-1.2*-mediated resistance against RKN *Meloidogyne javanica* (Atamian et al., 2012). We found that the *S. torvum* homologs of tomato WRKY31, 33, and 39 were specifically up-regulated upon infection with A2-O, but not with A2-J (Figure 2D). In the case of WRKY80, A2-J induced its expression at a later time point (3 DPI), while A2-O induced the WRKY80 homolog to much higher levels. These results suggest that A2-O, but not A2-J, strongly and specifically induces the expression of defense-related genes in *S. torvum*.

### Infection with A2-O Rapidly Induces the Expression of Sesquiterpene Synthases and Class III Peroxidases

Infection with A2-O caused an increase in expression of more genes and to a greater extent than A2-J (Figures 2B, 3A and Supplementary Figure 2). Importantly, at 1 DPI, A2-J did not induce any statistically significant changes in the expression of any genes, whereas A2-O induced 204 genes, suggesting that infection with A2-O rapidly induces the expression of early responsive genes, which is prevented or avoided in A2-J infection. Since the speed of a defense response is one of the most important factors for successful immunity against pathogens, we hypothesized there must be important defense components among the 204 up-regulated genes. We therefore performed a GO enrichment analysis to identify significantly represented GO terms amongst the 204 up-regulated genes (FDR ≤ 1E-04, Supplementary Table 5). The list of enriched GO terms was further reduced using “Reduce to most specific terms” option in Blast2GO to remove general GO terms and obtain only the most specific terms (Table 2). Some GO terms that were significantly enriched among the 204 genes were related to the biosynthesis of isoprenoids (“farnesyl diphosphate catabolic process (GO:0045339)”, “sesquiterpene biosynthetic process (GO:0051762)”, and “terpenoid biosynthetic process (GO:0016114)”). To follow up on this result, we checked the expression of all the genes up-regulated by A2-O that are related to isoprenoid biosynthesis and found that A2-O infection induced the expression of genes encoding sesquiterpene synthases, such as viridiflorene synthase, vetispiradiene synthase, germacrene C synthase-like protein, and 5-epiaristolochene synthase. Several other enzymes involved in isoprenoid biosynthesis, such as xanthoxin dehydrogenase-like protein and UDP-glycosyltransferase 91C1 were also up-regulated (Figure 3B). Sesquiterpene synthases convert farnesyl diphosphate to sesquiterpenes such as germacrene C, 5-epiaristolochene, viridiflorene, and vetispiradiene. Because some isoprenoids have nematicidal activity (Ohri and Pannu, 2009), it is possible that the sesquiterpenes produced by *S. torvum* in response to infection with A2-O are nematicidal and contribute to suppressing A2-O infection. Other GO terms significantly enriched among the 204 up-regulated genes were related to oxidative stress (“hydrogen peroxide catabolic process (GO:0042744)” and “response to oxidative stress (GO:0006979)”). The most up-regulated genes by A2-O in the GO term group were class III peroxidases, which are involved in lignification, cell elongation, seed germination, and response to abiotic and biotic stresses (Cosio and Dunand, 2009; Shigeto and Tsutsumi, 2016) (Supplementary Figure 3). The transcriptional up-regulation of the class III peroxidases is consistetnt with the fact that resistant tomato lines more strongly elevate peroxidase activity during RKN infection than susceptible lines (Zacheo et al., 1993).

**FIGURE 3.**
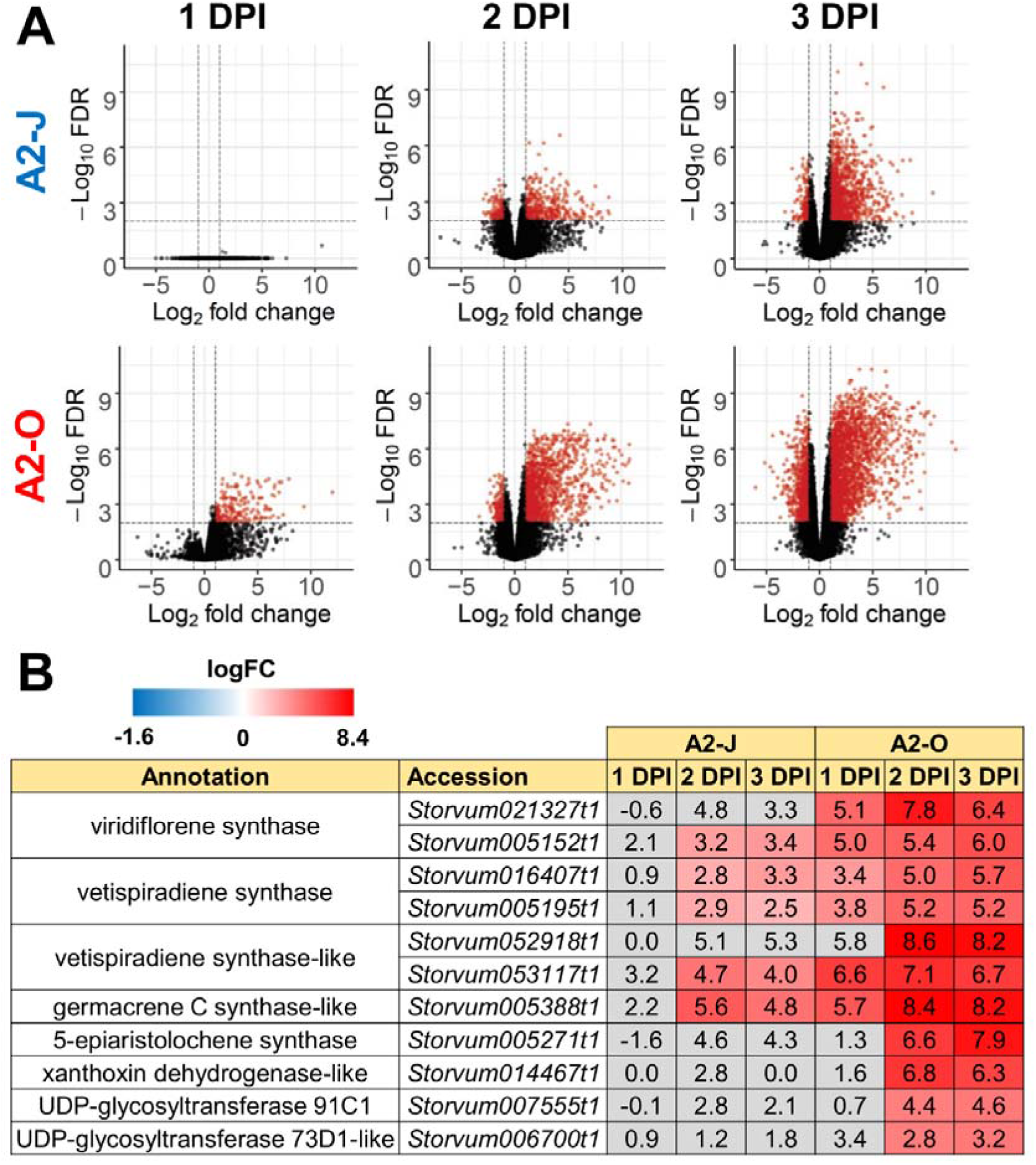
Infection with A2-O, but not with A2-J induced rapid expression of genes involved in *S. torvum* isoprenoid biosynthesis. **(A)** Volcano plots showing the distribution of differentially expressed genes in *S. torvum*infected with *M. arenaria* A2-J or A2-O compared to mock treatment. The logarithms of the fold change of individual genes are plotted against the negative logarithms of their FDR. Red dots represent genes up-regulated (logFC ≥ 1) or down-regulated (logFC ≤ −1) with an FDR < 0.01. (**B**) Infection with A2-O rapidly induced the expression of the genes involved in isoprenoid biosynthesis listed as “farnesyl diphosphate catabolic process (GO:0045339)”, “sesquiterpene biosynthetic process (GO:0051762)”, and “terpenoid biosynthetic process (GO:0016114)”. logFC values of the genes (compared to mock treatment) at 1, 2, and 3 DPI are shown. In the heatmap, grey boxes indicate no statistically significant difference at |logFC| ≥ 1 and FDR ≤ 0.01.

**TABLE 2.** Gene Ontology (GO) enrichment analysis of genes up-regulated by infection with *M. arenaria* A2-O at 1 DPI^a^

| GO term | Fold | FDR |
| --- | --- | --- |
| farnesyl diphosphate catabolic process (GO:0045339) | 347.4 | 7.9E-08 |
| sesquiterpene biosynthetic process (GO:0051762) | 248.2 | 4.0E-07 |
| hydrogen peroxide catabolic process (GO:0042744) | 23.6 | 6.6E-06 |
| response to oxidative stress (GO:0006979) | 11.8 | 2.2E-05 |
| terpenoid biosynthetic process (GO:0016114) | 21.9 | 7.1E-05 |

<sup>a</sup> The most specific GO terms enriched in 204 genes up-regulated by infection with A2-O at 1 DPI (FDR ≤ 1E-04) are listed with fold enrichment (Fold) and FDR values. “Reduce to most specific” option in Blast2GO was used to remove general GO terms. All of the significantly enriched GO terms are shown in Supplementary Table 5 (FDR ≤ 1E-04). Only GO category “Biological Process” is shown.

### Infection with A2-J Induces Genes Related to Cell Wall Modification and Transmembrane Transport

To identify the expression pattern of genes that are specific to A2-J or A2-O infection and common in both pathotypes, we clustered genes according to their transcript profiles by PCA with SOM clustering. SOM clustering grouped 6,502 genes into nine clusters based on their differential gene expression profiles after mock treatment or infection with either A2-J or A2-O (Figure 4 and Supplementary Table 6). 429 genes in Cluster 2 and 554 genes in Cluster 4 were specifically up-regulated after infection with A2-J. In contrast, 1,769 and 600 genes in Cluster 8 and 9, respectively, were specifically up-regulated after infection with A2-O. 1,000 genes in Cluster 7 were up-regulated after infection with either A2-J or A2-O (Figure 4B).

**FIGURE 4.**
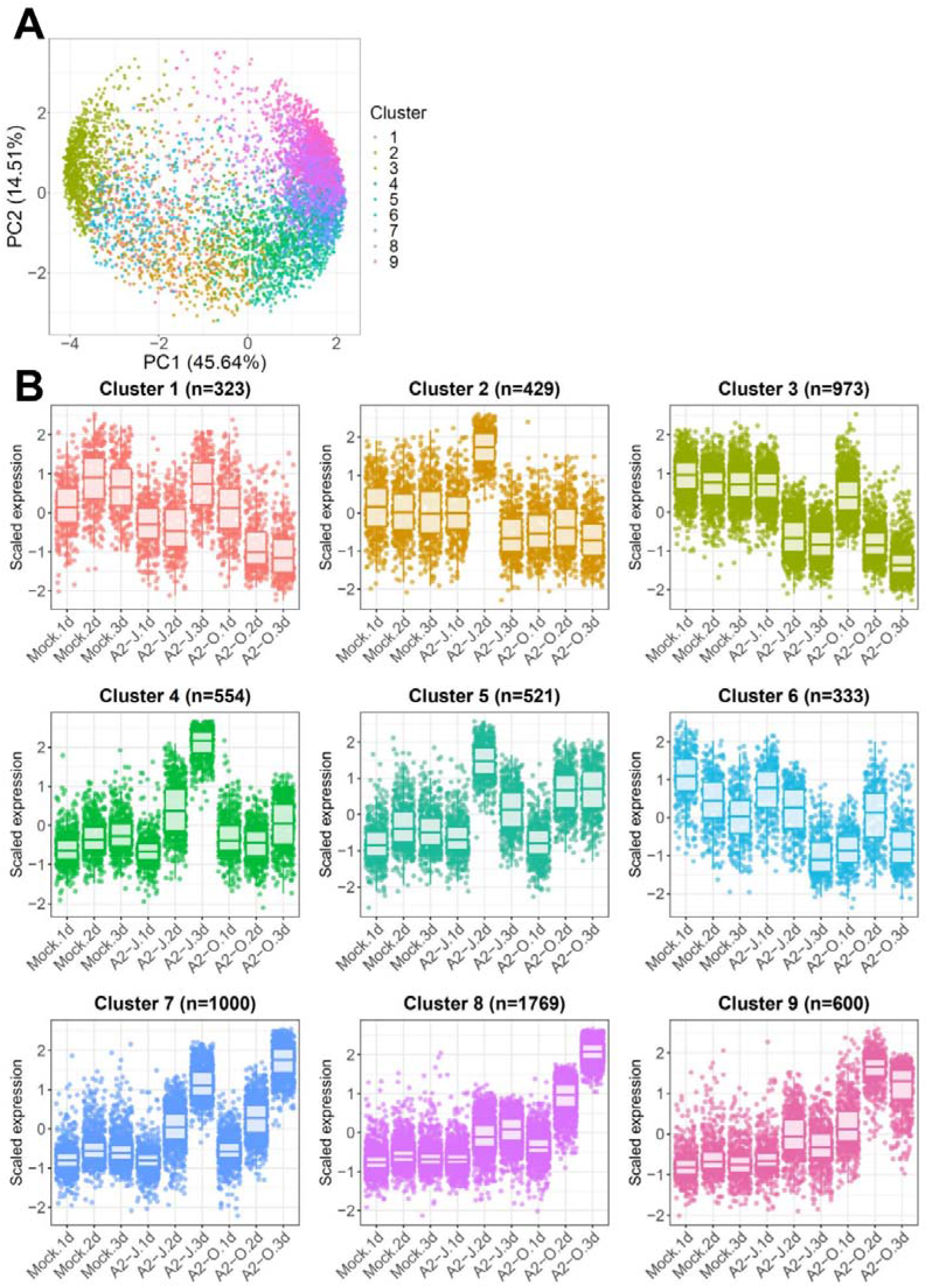
Principal component analysis (PCA) and self-organizing map (SOM) clustering of differentially expressed genes in *S. torvum*. (**A**) Clustering of gene expression patterns in *S. torvum* treated with DW (mock) or infected with A2-J or A2-O at 1, 2, and 3 DPI. The expression profiles of each gene were represented in a PCA space separated by PC1 and PC2, with SOM node memberships (cluster) represented by different colors and numbers. The percentage shown along the x or y-axis represents the percentage of variance explained by each component. A total of nine clusters with different expression patterns were defined. (**B**) Scaled expression profiles of genes by SOM clustering. The number of genes assigned to each SOM cluster is shown in parentheses. Each dot represents the value of the scaled expression of the gene. The upper and lower horizontal lines of the box correspond to the 25^th^ and 75^th^ percentiles. The horizontal bar in the box indicates the median. The upper and lower end of horizontal bars outside the box indicate 90^th^ and 10^th^ percentiles.

We once again used GO enrichment to identify functional terms enriched in the genes in each cluster (FDR ≤ 1E-04, Supplementary Table 7) by further filtering enriched GO terms using the “Reduce to most specific terms” option in Blast2GO. In Cluster 4 (specifically A2-J up-regulated genes), significantly enriched GO terms were related to cell wall remodeling, including “cell wall modification (GO:0042545)”, “cell wall organization or biogenesis (GO:0071554)”, and “pectin catabolic process (GO:0045490)” (Table 3 and Supplementary Table 7). This is consistent with the observation that the expansion of giant cells is associated with an increase in cell wall thickness (Bartlem et al., 2014; Bozbuga et al., 2018). GO terms associated with the significantly A2-J up-regulated genes include enzymes such as cellulose synthase-like protein, xyloglucan endotransglucosylase/hydrolase protein, and a non-catalytic subunit of a polygalacturonase isozyme (Supplementary Figure 4A). The transcriptional up-regulation of these enzymes is consistent with the presence of the common polysaccharides pectic homogalacturonan, xyloglucan, and pectic arabinan in the cell walls of giant cells (Bozbuga et al., 2018). Similarly, A2-J infection also activated the expression of COBRA-like protein, expansin, and LRR-RLK PXC1, which play important roles in cellulose deposition, loosening of cell walls, and secondary wall formation (Cosgrove, 2000; Brown et al., 2005; Wang et al., 2013; Kumar et al., 2016) (Supplementary Figure 4A). Another GO term significantly enriched in Cluster 4 was “transmembrane transport (GO:0055085)” (Supplementary Table 7). Other significantly up-regulated genes encode sugar transporter ERD6-like protein (Hammes et al., 2005) and amino acid transporter family protein (Elashry et al., 2013). These transporters may promote the uptake of nutrients into giant cells or alter transportation through cells surrounding giant cells (Supplementary Figure 4B). We also performed GO enrichment analyses for Cluster 2, but no GO terms were enriched.

**TABLE 3.** Gene Ontology (GO) enrichment analysis of Cluster 4 genes up-regulated specifically by infection with *M. arenaria* A2-J^a^

| Cluster | GO term | Fold | FDR |
| --- | --- | --- | --- |
| Cluster 4 | pectin catabolic process (GO:0045490) | 16.9 | 8.1E-07 |
|  | cell wall modification (GO:0042545) | 14.7 | 8.1E-05 |
|  | negative regulation of catalytic activity (GO:0043086) | 7.0 | 9.8E-05 |

<sup>a</sup> The most specific GO terms enriched in Cluster 4 ( $\text{FDR} \leq 1\text{E-}04$ ) are listed with fold enrichment (Fold) and FDR values. “Reduce to most specific” option in Blast2GO was used to remove general GO terms. All of the significantly enriched GO terms in each cluster are shown in Supplementary Table 7 ( $\text{FDR} \leq 1\text{E-}04$ ). Only GO category “Biological Process” is shown.

In addition to GO enrichment analyses, we looked for interesting genes whose expression were dramatically up-regulated in Clusters 2 and 4. These clusters included genes encoding chalcone synthase and a spermidine synthase that were specifically and highly expressed after infection with A2-J (Figure 5A). Chalcone synthase is the first enzyme of the flavonoid biosynthetic pathway (Dao et al., 2011), and spermidine synthase is a key enzyme involved in polyamine biosynthesis (Liu et al., 2007). In summary, A2-J infection significantly and specifically up-regulates genes related to cell wall modification and membrane transport, chalcone synthase, and spermidine synthase at an early phase of gall formation.

**FIGURE 5.**
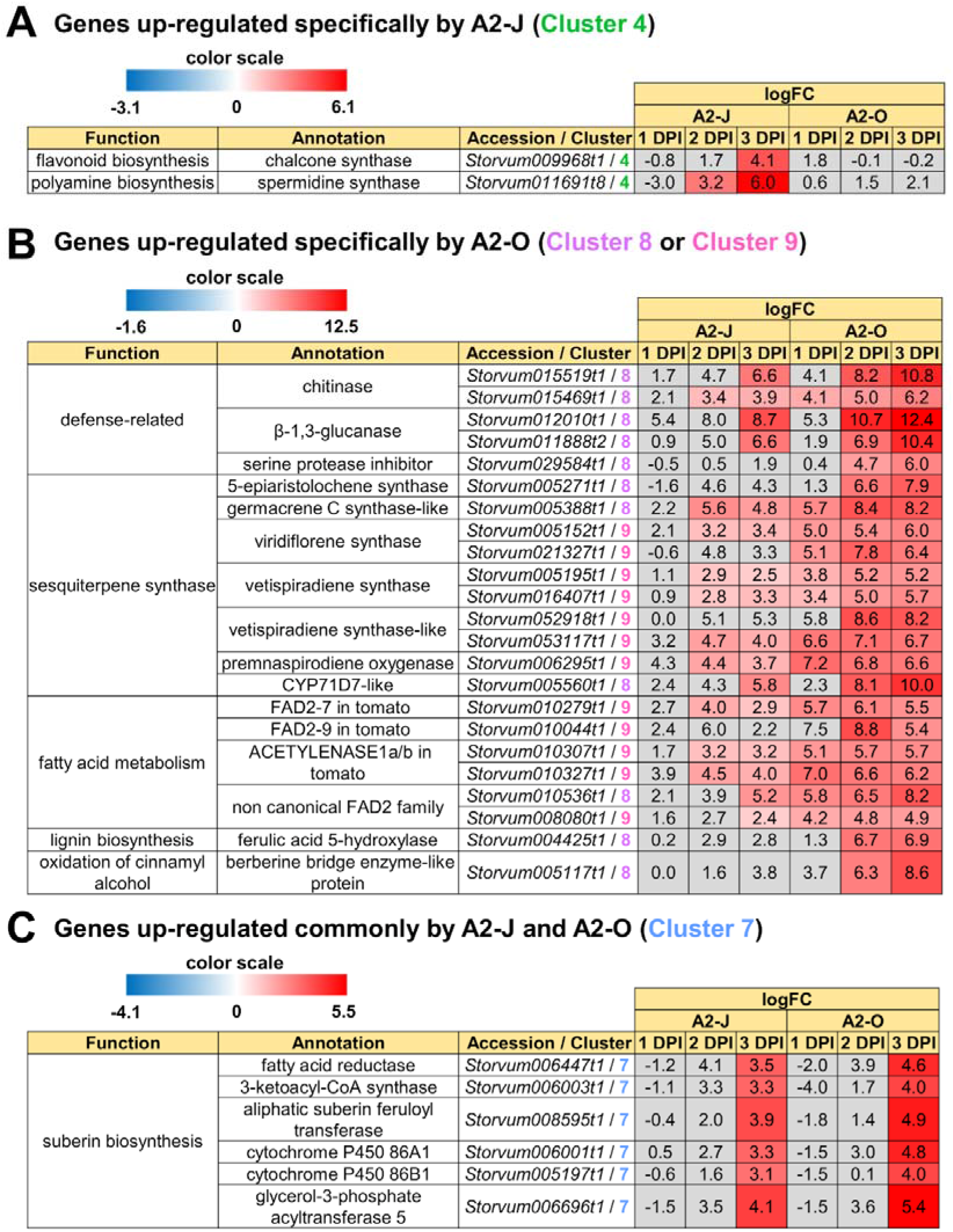
Up-regulated genes in Clusters 4, 7, 8, and 9. Highly up-regulated genes (logFC ≥ 4, FDR ≤ 0.01) in each cluster are listed. (**A**) Infection with A2-J specifically up-regulated the expression of chalcone synthase and spermidine synthase (Cluster 4). (**B**) Infection with A2-O dramatically up-regulated defense-related proteins, sesquiterpene synthase, fatty acid desaturase 2 (FAD2), ferulic acid 5-hydroxylase, and berberine bridge enzyme-like protein (Cluster 8 and 9). (**C**) Both A2-J and A2-O induced the expression of suberin biosynthetic enzymes (Cluster 7). logFC values of the genes compared to mock treatment at 1, 2, and 3 DPI are shown. The heatmap grey to indicate values with no statistically significant differences at |logFC| ≥ 1 and FDR ≤ 0.01.

### Infection with A2-O Induces Genes Related to Defense Responses

In Cluster 8 (genes specifically up-regulated by A2-O), GO terms that were significantly enriched were related to defense responses, including “defense response to fungus (GO: 0050832)”, “defense response to bacterium (GO: 0042742)”, “killing of cells of other organism (GO:0031640)”, and “regulation of salicylic acid biosynthetic process (GO: 0080142)”. In addition, GO terms involved in lignin biosynthesis, including “lignin biosynthetic process (GO:0009809)” was also overrepresented in Cluster 8 (Table 4 and Supplementary Table 7). In Cluster 9, the significantly enriched GO terms were related to biosynthesis of isoprenoids (“sesquiterpene biosynthetic process (GO: 0051762)”, “terpenoid biosynthetic process (GO: 0016114)”, and “farnesyl diphosphate catabolic process (GO:0045339)”) (Table 4 and Supplementary Table 7).

**TABLE 4.** Gene Ontology (GO) enrichment analysis of Cluster 8 and 9 genes up-regulated specifically by infection with *M. arenaria* A2-O^a^

| Cluster | GO term | Fold | FDR |
| --- | --- | --- | --- |
| Cluster 8 | glutathione metabolic process (GO:0006749) | 18.1 | 1.8E-27 |
|  | recognition of pollen (GO:0048544) | 8.9 | 1.2E-13 |
|  | protein ubiquitination (GO:0016567) | 3.5 | 1.2E-11 |
|  | defense response to fungus (GO:0050832) | 8.2 | 5.4E-10 |
|  | cellular oxidant detoxification (GO:0098869) | 4.9 | 6.1E-09 |
|  | defense response to bacterium (GO:0042742) | 6.3 | 1.2E-08 |
|  | hydrogen peroxide catabolic process (GO:0042744) | 6.6 | 5.6E-08 |
|  | lignin biosynthetic process (GO:0009809) | 11.7 | 9.4E-08 |
|  | protein autophosphorylation (GO:0046777) | 4.1 | 1.0E-07 |
|  | chitin catabolic process (GO:0006032) | 18.2 | 3.1E-07 |
|  | response to oxidative stress (GO:0006979) | 3.6 | 3.5E-06 |
|  | mRNA transcription (GO:0009299) | 21.5 | 7.1E-06 |
|  | killing of cells of other organism (GO:0031640) | 18.8 | 1.7E-05 |
|  | regulation of salicylic acid biosynthetic process (GO:0080142) | 15.8 | 5.3E-05 |
|  | response to abscisic acid (GO:0009737) | 3.7 | 9.1E-05 |
| Cluster 9 | negative regulation of endopeptidase activity (GO:0010951) | 17.7 | 5.9E-07 |
|  | farnesyl diphosphate catabolic process (GO:0045339) | 118.1 | 2.0E-06 |
|  | sesquiterpene biosynthetic process (GO:0051762) | 84.4 | 1.1E-05 |
|  | terpenoid biosynthetic process (GO:0016114) | 11.2 | 1.9E-05 |

<sup>a</sup> The most specific GO terms enriched in Cluster 8 and 9 ( $FDR \leq 1E-04$ ) are listed with fold enrichment (Fold) and FDR values. “Reduce to most specific” option in Blast2GO was used to remove general GO terms. All of the significantly enriched GO terms in each cluster are shown in Supplementary Table 7 ( $FDR \leq 1E-04$ ). Only GO category “Biological Process” is shown.

We also found that the genes that are highly expressed after infection with A2-O in Cluster 8 and 9 include (1) defense-related genes encoding chitinase, β-1,3-glucanase, and serine protease inhibitor, (2) sesquiterpene synthase, (3) fatty acid desaturase 2 (FAD2), (4) ferulic acid 5-hydroxylase (F5H) which is involved in lignin biosynthesis, (5) berberine bridge enzyme (BBE)-like protein, which is involved in oxidation of cinnamyl alcohol (Figure 5B). Fatty acids are major and essential components of all plant cells and are also precursors for a variety of plant metabolites, including signaling molecules and phytoalexins (Ohlrogge and Browse, 1995; Lim et al., 2017). *FAD2* encodes Δ12-desaturase that catalyzes the conversion of oleic acid (C18:1) to linoleic acid (C18:2) (Ohlrogge and Browse, 1995). The Arabidopsis genome has only a single *FAD2* gene (AT3G12120), but most other plant species carry multiple *FAD2* homologs (Cao et al., 2013; Lee et al., 2020). The duplication of *FAD2* genes in plants would have enabled the functional diversification of these enzymes, leading to divergent catalytic activities and the synthesis of novel metabolites. For example, recent studies have shown that tomato has non-canonical FAD2 family proteins that lack Δ12-desaturase activity (Jeon et al., 2020; Lee et al., 2020). In particular, ACET1a/b (Solyc12g100240 and Solyc12g100260) and FAD2-9 (Solyc12g100250) are non-canonical FAD2 involved in the biosynthesis pathway from linoleic acid to a phytoalexin, falcarindiol (Jeon et al., 2020). Falcarindiol has not only anti-bacterial and anti-fungal activities but also nematicidal activity to *M. incognita* and pinewood nematode *Bursaphelenchus xylophilus* (Liu et al., 2016). Infection with A2-O rapidly induced the expression of *ACET1a/b* and *FAD2-9* (Figure 5B), suggesting that infection with A2-O rapidly activates a biosynthesis pathway similar to the falcarindiol pathway, but the production of falcarindiol by *S. torvum* needs to be experimentally confirmed in the future.

### Infection with A2-O Induces Lignin Accumulation in *S. torvum*

The GO enrichment analysis of Cluster 8 revealed that “lignin biosynthetic process (GO:0009809)” was significantly enriched (Table 4 and Supplementary Table 7), and that the expression of *F5H* in Cluster 8 was very high and specifically induced after infection with A2-O (Figure 5B). These results suggest that the infection with A2-O transcriptionally activates lignin biosynthesis. Lignin is a phenylpropanoid polymer that is deposited predominantly in the secondary cell wall, making the cell wall rigid and impervious to water (Vanholme et al., 2010). Lignin polymer is synthesized via oxidative combinational coupling of lignin monomers (or monolignols), namely *p*-coumaryl alcohol, sinapyl alcohol and coniferyl alcohol. The lignin subunits constituted by these monolignols are *p*-hydroxyphenyl (H), syringyl (S), and guaiacyl (G) groups, respectively. All of the monolignols are synthesized from phenylalanine through the general phenylpropanoid and monolignol-specific pathways (Vanholme et al., 2012) (Figure 6A). Normally, lignin deposition occurs in the root endodermis of the differentiation zone and constitutes the Casparian strip, which functions as a physical barrier that prevents free diffusion of solutes and ions between the xylem and the soil (Robbins 2nd et al., 2014). However, biosynthesis and deposition of lignin can be induced in response to biotic stresses (Miedes et al., 2014; Mutuku et al., 2019), which prompted a closer examination of the expression patterns of genes involved in the lignin biosynthetic pathway whose expression was up-regulated by infection with either A2-J or A2-O (logFC ≥ 1, FDR ≤ 0.01) (Figure 6B). Infection with A2-O induced the expression of genes encoding phenylalanine ammonia-lyase (PAL), cinnamate 4-hydroxylase (C4H), 4-coumaroyl-CoA ligase (4CL), *p*-hydroxycinnamoyl-CoA:shikimate *p*-hydroxycinnamoyl transferase (HCT), caffeoyl-CoA O-methyltransferase (CCoAOMT), cinnamoyl-CoA reductase (CCR), F5H, caffeic acid O-methyltransferase (COMT), and cinnamyl alcohol dehydrogenase (CAD). Infection with A2-J also induced the expression of some of these genes, but to a much lesser extent than A2-O (Figure 6B). Phloroglucinol staining of infected roots allowed us to visualize the intensity and location of lignin accumulation (Figure 7). Infection with A2-O, but not with mock treatment, induced ectopic accumulation of lignin in root tips. With A2-J infection, the area of the root proximal to gall tissue was very slightly stained with phloroglucinol, but the gall itself had little or no detectable phloroglucinol staining. These differences in lignin staining intensity may reflect differences in the expression of lignin biosynthetic genes after infection with A2-J and A2-O (Figure 5B).

**FIGURE 6.**
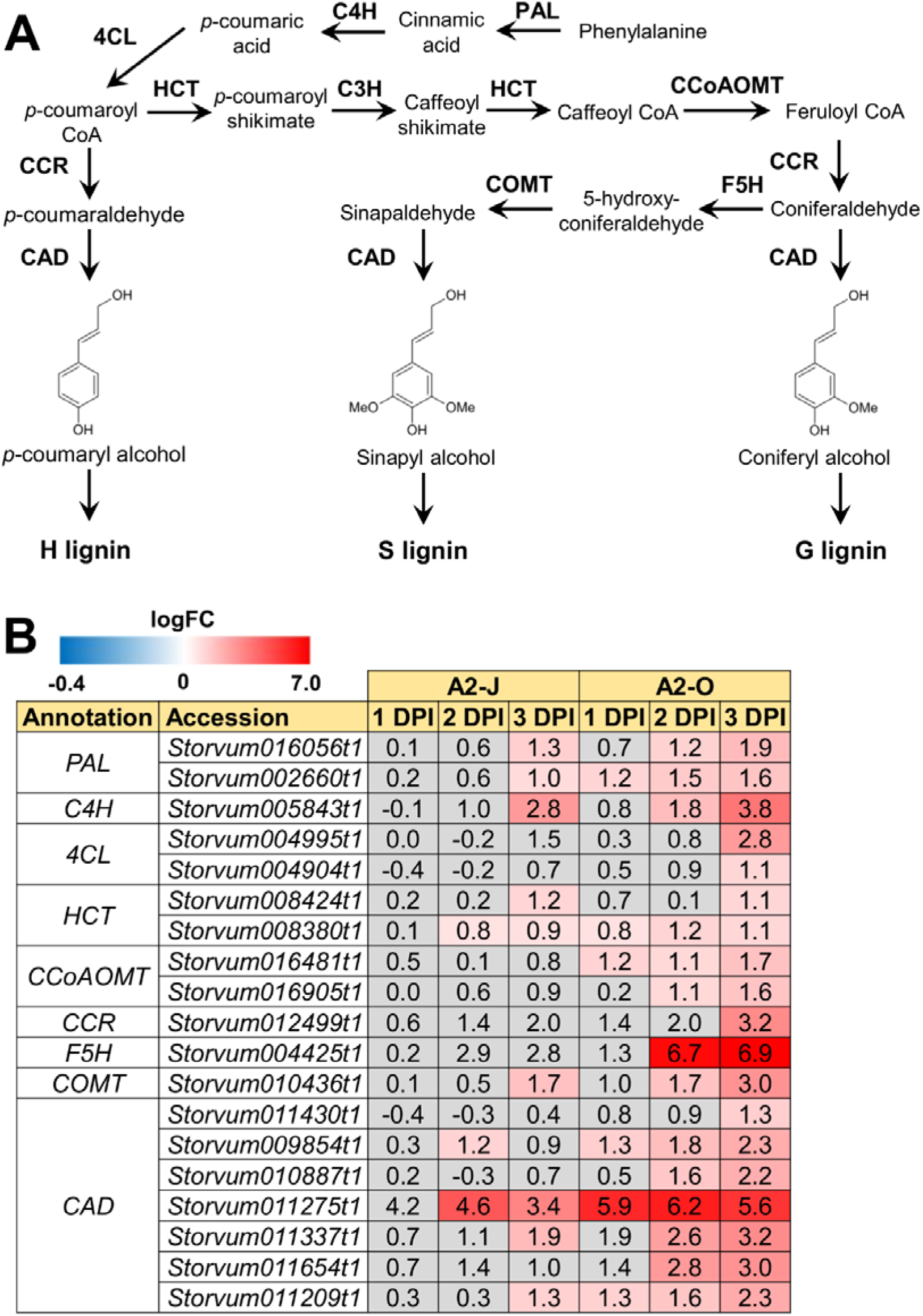
Infection with A2-O induced the expression of lignin biosynthetic genes. (**A**) Overview of the lignin biosynthesis pathway. PAL, phenylalanine ammonia-lyase; C4H, cinnamate 4-hydroxylase; 4CL, 4-coumaroyl-CoA ligase; HCT, p-hydroxycinnamoyl-CoA:shikimate p-hydroxycinnamoyl transferase; C3H, p-coumarate 3-hydroxylase; CCoAOMT, caffeoyl-CoA O-methyltransferase; CCR, cinnamoyl-CoA reductase; F5H, ferulate 5-hydroxylase; COMT, caffeic acid O-methyltransferase; CAD, cinnamyl alcohol dehydrogenase. (**B**) Expression patterns of genes involved in the lignin biosynthetic pathway whose expression was significantly up-regulated by infection with A2-J or A2-O (logFC ≥ 1, FDR ≤ 0.01). logFC values of the genes compared to mock treatment at 1, 2, and 3 DPI are shown. The heatmap uses grey to indicate values with no statistically significant differences at |logFC| ≥ 1 and FDR ≤ 0.01.

### Infection with A2-O or A2-J Induces the Expression of Genes Related to Suberin Biosynthesis

There was no enrichment of specific GO terms in Cluster 7 (up-regulated by the infection with A2-J or A2-O) (Supplementary Table 7). However, we found that both A2-J and A2-O strongly activate the expression of suberin biosynthetic genes, including *aliphatic suberin feruloyl transferase (ASFT), cytochrome P450 86A1 (CYP86A1), cytochrome P450 86B1 (CYP86B1), glycerol-3-phosphate acyltransferase 5 (GPAT5)*, and *β□ketoacyl□CoA synthase (KCS)* (Figure 5C), but A2-O induced slightly higher expression of these genes than A2-J. Suberin is a cell wall component that restricts water loss, nutrient elution, and pathogen infection (Dean and Kolattukudy, 1976; Lulai et al., 2008; Leide et al., 2012; Froschel et al., 2020). It is normally deposited in the cell walls of endodermal cells, but not in the root tip (Barberon, 2017), and several reports showed that suberin synthesis is induced in wounded tissues (Dean and Kolattukudy, 1976; Bernards and Lewis, 1998). Considering that both A2-J and A2-O induced the expression of suberin biosynthetic genes, the expression of these genes was possibly up-regulated by a generalized wounding signal during nematode infection. We therefore investigated the expression levels of the wound-responsive genes *ATAF2*, which encodes a wound-responsive NAC transcription factor (Collinge and Boller, 2001; Wang and Culver, 2012), and *pathogen-related 4* (Stanford et al., 1989; Marhavý et al., 2019) (Supplementary Figure 5). Not surprisingly, the expression patterns of wound-responsive genes were similar to those of the suberin biosynthetic genes. These results suggest that infection with A2-O may cause more wounding and damage than A2-J, but that the wounding signal generated during the earliest events in nematode infection, whether a resistant or susceptible reaction occurs later, may induce the expression of suberin biosynthetic genes.

Considering the suberin biosynthetic gene transcription data, we predicted that infection with A2-O or A2-J would induce the accumulation of suberin. To test our prediction, we examined and measured the chemical composition of aliphatic suberin at the nematode infection site, but total aliphatic suberin content did not increase after A2-J or A2-O infection (Welch’ s *t*-test with Bonferroni correction, *p*≤ 0.05) (Figure 8A). However, the abundance of single monomers, ω-hydroxy acid (ω-OH-acid) C16 and α,ω-diacid C16, were significantly higher after infection with A2-O than mock treatment (*p*≤ 0.05, Welch’s *t*-test followed by Bonferroni correction) (Figure 8B). The large variation in the data may be due to variation in the number of infected root tips, nematode aggressiveness, and/or the severity of wounding, which can vary by experiment and cause disparity in the proportions of suberized areas among the fragile root tips. Because infection with RKN *M. incognita* induces the expression of suberin biosynthetic genes and a patchy suberin accumulation pattern within the cell layer surrounding the infection site in later stages of infection (10 DPI) in susceptible Arabidopsis plants (Holbein et al., 2019), it is possible that the synthesis of large amounts of suberin takes longer in *S. torvum* that what was allowed for in this assay.

## DISCUSSION

### Comparative Transcriptome Analyses for Host Defense and Susceptible Responses at the Early Stages of Nematode Infection

In this study we established an experimental system using a single cultivar of *S. torvum* and two pathotypes of RKNs *M. arenaria* A2-J, which is able to infect and establish a parasitic relationship with the plant, and A2-O, which induces a resistance response. To the best of our knowledge, this is the first comparative RNA-seq analyses in a single cultivar with virulent or avirulent nematodes. Using this experimental system, we were able to catalog changes in gene expression at the very early stage of infection (1-3 DPI) with a high degree of sensitivity compared to previous studies (Bagnaresi et al., 2013; Postnikova et al., 2015; Xing et al., 2017; Shukla et al., 2018). Because there are clear morphological differences in root tips infected with A2-J and A2-O within 4 DPI, the sum of host responses by this stage of infection likely has determined the outcome of infection. Interestingly, at 1 DPI, A2-O induced the expression of genes encoding class III peroxidases, fatty acid desaturases, and enzymes involved in isoprenoid biosynthesis (Figure 3B; Figure 5B; Supplemental Figure 3), whereas A2-J had not induced any statistically significant changes in the expression of defense response genes (Figure 3A). These results suggest that A2-J initially evades recognition by *S. torvum* and/or actively suppresses the induction of transcriptional changes in the host. Our results also show that A2-J induces the expression of genes associated with susceptible responses that are related to gall formation at around 2-3 DPI (Supplementary Figure 4). The delay in expression of susceptibility-associated genes is consistent with the length of time it takes for nematodes to migrate through the plant’s vascular system to their target cells.

Sampling only infected root tips reduced background plant transcripts and enabled us to detect dynamic changes in gene expression that are directly associated with the establishment of a parasitic relationship or with active host defense. The range of calculated logFC values of root tip genes expressed at 3 DPI with A2-O was from −8 to 12.8 (that is, from 1/256 to 7131 times), which is quite broad for very early stages of infection. For example, in previous transcriptome analyses using *S. torvum* during infection with an avirulent isolate of *M. incognita*, chitinase expression in whole roots was only 2-4 times higher upon infection than in uninfected roots (Bagnaresi et al., 2013). In contrast, our analyses showed that the expression of chitinases in the root tips infected with A2-O at 3 DPI was several hundred times higher than in the mock treatment, clearly demonstrating the high sensitivity of this method for host-parasite transcriptomics (Figure 5B). This level of sensitivity allowed us to identify genes that had not previously been connected with resistance or susceptibility in this system. In addition, comparative transcriptome analyses of *S. torvum* combined with SOM clustering can be used to pinpoint genes that are associated with susceptibility or resistance.

### Infection with A2-J Induces the Expression of Genes Related to Cell Wall Modification and Spermidine Synthase

A2-J specifically induced the expression of cell wall modification enzymes such as cellulose synthase-like protein, xyloglucan endotransglucosylase/hydrolase protein, and a non-catalytic subunit of the polygalacturonase isozyme (Table 3 and Supplementary Figure 3A). The cell walls of giant cells require thickening and loosening to allow cell expansion that increases the surface area of the plasma membrane, and to support nutrient uptake by the nematodes (Bartlem et al., 2014; Bozbuga et al., 2018; Meidani et al., 2019). The cell walls of giant cells and syncytial feeding sites induced by CNs contain high-ester pectic homogalacturonan, xyloglucan, and pectic arabinan (Davies et al., 2012; Zhang et al., 2017b), suggesting that these polysaccharides are responsible for the flexible properties of feeding site cell walls. Our transcriptome results confirm the importance of cell wall modification enzymes during gall formation.

In addition to cell wall modification enzymes, infection with A2-J specifically induced the expression of spermidine synthase (Figure 5A). Interestingly, a virulence effector protein secreted from CNs, 10A06, functions through its interaction with Arabidopsis spermidine synthase 2 (Hewezi et al., 2010) by increasing spermidine concentrations, subsequently increasing polyamine oxidase activities. An increase in polyamine oxidase activity results in the induction of cellular antioxidant machinery in syncytia and disruption of SA-mediated defense signaling. Although there is no clear homolog of 10A06 in RKNs such as *M. incongnita*, it is possible that there is a functional ortholog of 10A06 that up-regulates the expression of spermidine synthase. Potentially, transcriptional activation of the spermidine synthase gene by PPNs may turn out to be a common strategy for suppressing plant immunity.

### Infection with A2-O Up-regulates Genes Related to Sesquiterpene Biosynthesis and Fatty Acid Metabolism

Infection with A2-O, but not with A2-J, strongly induced genes encoding sesquiterpene synthases (Table 4 and Figure 5B). High levels of expression of these genes occurs at the very early stages of infection (1 DPI), which sets these genes apart from the other genes induced by A2-O infection (Figure 3B and Figure 5B), suggesting the importance of sesquiterpenes as an early line of defense against PPNs. Capsidiol, a sesquiterpene, is the major phytoalexin produced in the Solanaceae plants *Nicotiana* spp. and *Capsicum* spp. in response to fungal and bacterial infection (Grosskinsky et al., 2011; Song et al., 2019). Capsidiol is toxic to many oospore and fungal pathogens, such as *Phytophthora capsici* and *Botrytis cinerea* (Stoessl et al., 1972; Ward et al., 1974), and suppresses the mobility of false root-knot nematode *Nacobbus aberrans* (Godinez-Vidal et al., 2010). We found that A2-O induces the expression of CYP71D7-like protein, the closest homologue to CYP71D20 from *Nicotiana benthamiana*,which converts 5-epiaristolochene to capsidiol (Ralston et al., 2001). A2-O also induces the expression of genes encoding 5-epiaristolochene synthase, which converts farnesyl diphosphate to 5-epiaristolochene (Vogeli and Chappell, 1988; Facchini and Chappell, 1992). Although capsidiol production is not common in *Solanum* spp., *S. torvum* may produce similar sesquiterpene derivatives that are toxic to PPNs. Another sesquiterpene that may be involved in resistance is the phytoalexin solavetivone, because A2-O induces the expression of genes encoding vetispiradiene synthase and premnaspirodiene oxygenase, which sequentially convert farnesyl diphosphate to solavetivone via vetispiradiene (Takahashi et al., 2007). Although the nematicidal activity of solavetivone has not yet been reported, genetic analyses indicate that the production of solavetivone is associated with resistance in potato against CN *Globodera rostochiensis* (Desjardins et al., 1997). It would be interesting to test the nematicidal activity of these sesquiterpenes and their production after A2-O infection in *S. torvum*.

We also found that infection with A2-O induces the expression of genes encoding the non-canonical FAD2 proteins ACET1a/b (Solyc12g100240 and Solyc12g100260) and FAD2-9 (Solyc12g100250), which are involved in the biosynthesis of falcarindiol, a modified fatty acid found in a variety of plants, including Solanaceae. It is toxic to RKNs and pinewood nematodes (Liu et al., 2016), but it is not known if falcarindiol is produced by *S. torvum*. In any case, the higher expression of FAD2 after A2-O infection suggests the importance of fatty acid metabolism in plant immunity against PPNs. We consistently found that A2-O infection results in the accumulation of linoleic acid (C18:2), while infection with A2-J reduced linoleic acid (C18:2) and palmitic acid (C16:0) (Supplementary Figure 6). Increases in linoleic acid elicits resistance to the fungal pathogen *Colletotrichum gloeosporioides* in avocado (Madi et al., 2003), and to *B. cinerea* in bean plants (Ongena et al., 2004). The importance of fatty acids in resistance is further supported by the fact that PPNs secrete fatty-acid- and retinol-binding family proteins as a mechanism to increase susceptibility (Prior et al., 2001; Iberkleid et al., 2013; Iberkleid et al., 2015). Thus, the battle over fatty acid synthesis is likely to be important in *S. torvum*-PPN interactions.

### Lignin Accumulation as a Defense against PPNs

Because PPNs penetrate the cell wall and migrate within roots, reinforcement of cell walls by lignin accumulation has been implicated as an effective defense response to PPNs (Holbein et al., 2016; Sato et al., 2019). In fact, several studies showed that PPN infection induces more extensive lignin accumulation in resistant plants than in susceptible plants. For example, Veronico et al. (2018) performed a histochemical analysis of lignin in the roots of susceptible and resistant tomatoes infected with *M. incognita*, and found that accumulation of higher lignin levels in the root tissues (e.g. cortical cells) of resistant tomato than in susceptible varieties. Similarly, upon infection with the cereal CN *Heterodera avenae*, a resistant wheat cultivar gives a strong lignin accumulation response in the walls of cells affected by nematode infection, but a susceptible cultivar responded with only minor additional lignification, and only in cell walls mechanically damaged during nematode penetration (Andres et al., 2001). Since these studies used resistant and susceptible lines, it is certainly possible that differences in lignin accumulation are due to multiple differences in genetic background, and thus there would not necessarily be a direct relationship to resistance. In the present study, by using a single host cultivar with virulent and avirulent nematodes, we showed that susceptible and resistant responses are correlated with the levels of lignin. Infection with A2-O significantly up-regulated genes involved in lignin biosynthesis, and induced lignin accumulation near the root tip (Figure 6B and Figure 7). In contrast, infection with A2-J induced lignin biosynthetic genes to some limited extent that did not result in additional lignin accumulation. We also found that infection with A2-O significantly up-regulated the expression of class III peroxidases, some of which are thought to be involved in the polymerization of lignin (Zacheo et al., 1993; Zacheo et al., 1997; Marjamaa et al., 2009). The importance of lignin accumulation in PPN resistance is also supported by the fact that plant defense inducers such as β-aminobutyric acid, thiamine, sclareol, and benzothiadiazole induce lignin biosynthesis and enhance resistance against PPNs (Fujimoto et al., 2015; Ji et al., 2015; Huang et al., 2016; Veronico et al., 2018). Thus, ectopic lignification may result in cell wall reinforcement that restricts RKN invasion and migration.

**FIGURE 7.**
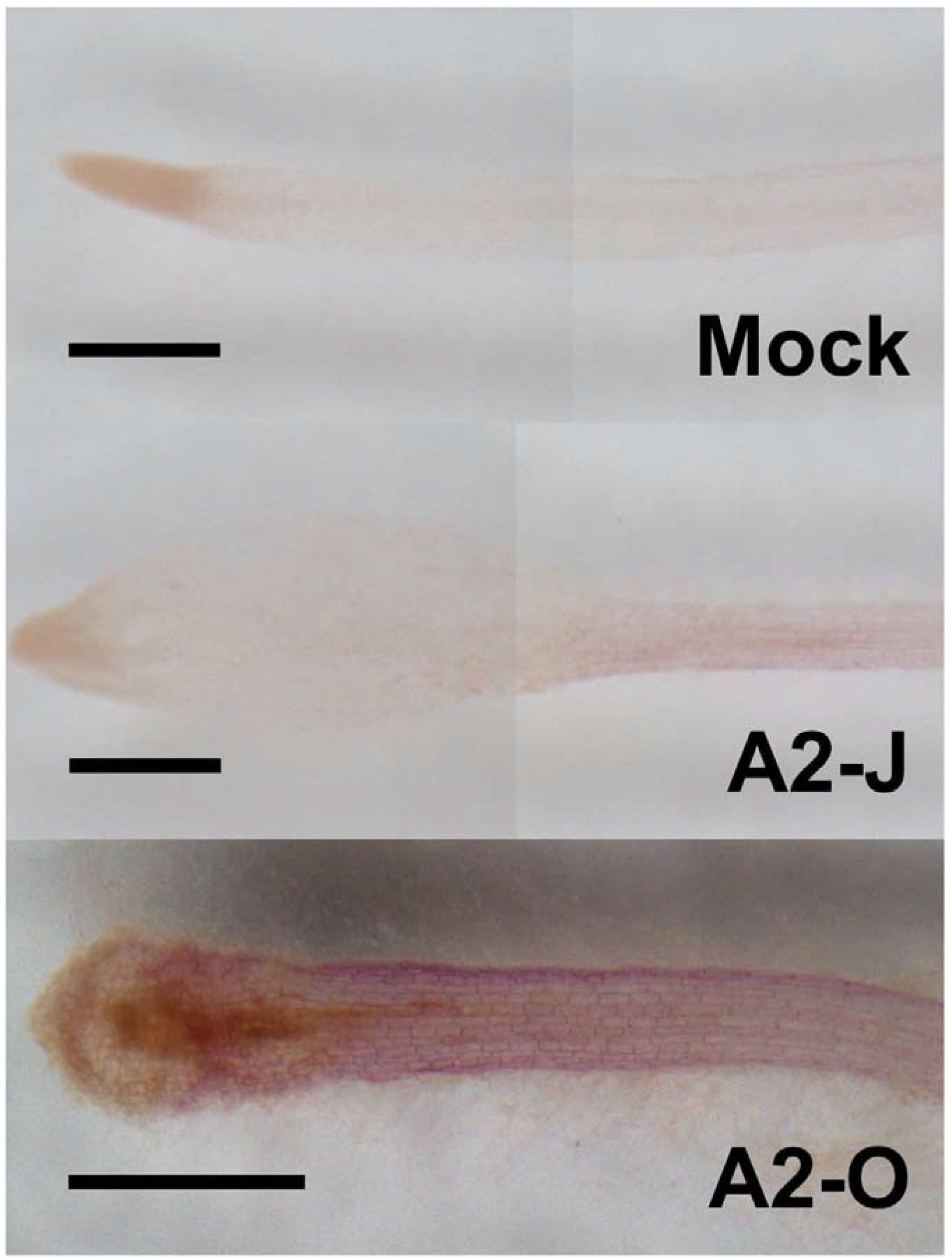
Infection with A2-O but not with A2-J induced lignin deposition. Phloroglucinol staining of lignin (red) in *S. torvum* root tips treated with SDW (mock) or infected with A2-J or A2-O for three days. Scale bars indicate 0.5 mm. This experiment was performed four times with the same results.

In contrast to the resistance interaction with A2-O, infection with A2-J did not induce lignin accumulation. One potential mechanism to explain this is that A2-J may alter the metabolic flux of lignin biosynthesis by specifically inducing the expression of the gene encoding chalcone synthase, which converts *p*-coumaroyl-CoA into naringenin chalcone (Figure 5A). The biosynthetic pathways leading to lignin and flavonoids diverge at the common intermediate *p*-coumaroyl CoA, so higher chalcone synthase expression should decrease lignin content. Indeed, silencing of the chalcone synthase gene increases lignin deposition in wheat (Eloy et al., 2017).

Activation of the phenylpropanoid biosynthesis pathway may also be involved in the production of anti-PPN secondary metabolites. For example, *trans*-cinnamaldehyde is a phenylpropanoid compound that is highly toxic to RKNs and pinewood nematodes (Oka, 2001; Kong et al., 2007). Cinnamaldehyde can be synthesized from L-phenylalanine, and its biosynthesis requires enzymatic reactions mediated by PAL, 4CL, and CCR (Bang et al., 2016). Expression of genes encoding these enzymes was specifically induced in A2-O-infected *S. torvum* (Figure 6B). We also observed a drastic induction of a BBE-like protein homologous to AtBBE-like 13 and AtBBE-like 15 (Figure 5B), which can convert cinnamyl alcohol to cinnamaldehyde in Arabidopsis (Daniel et al., 2015). Thus, *S. torvum* may activate the production of cinnamaldehyde to suppress PPN infection.

### Both A2-J and A2-O Induce the Expression of Suberin Synthetic Genes

Suberin is a lipid-phenolic biopolyester deposited in the cell walls of the root endodermis. Suberin lamellae and the lignin-based Casparian strip in the endodermis form the apoplastic diffusion barrier between vascular tissues and outer ground tissues to enforce selective absorption of water and nutrients (Alassimone et al., 2012; Vishwanath et al., 2015). The suberized endodermis may also a play role in defense against pathogens as the last line of defense before *pathogens* invade the vasculature (Thomas et al., 2007). Moreover, several reports showed that fungal or viral attack induces the deposition of suberin in the walls of cells in and around the site of penetration (Tippett and Hill, 1984; Kolattukudy and Espelie, 1989). In contrast, the virulent vascular pathogen, *Verticillium longisporum* reduces suberin deposition by suppressing the translation of genes involved in suberin biosynthesis (Froschel et al., 2020). These results may indicate the general importance of suberin in plant immunity. Infection with A2-J or A2-O induced the expression of genes involved in suberin biosynthesis at the root tip, with A2-O induction being slightly higher (Figure 5C). Although there was no statistical difference in total aliphatic suberin content, infection with A2-O increased the accumulation of suberin monomers such as ω-OH-acid C16 and α, ω-diacid C16 (Figure 8). These results suggest that *de novo* suberin synthesis is induced at the site of infection. Considering that A2-J also induces the expression of suberin biosynthesis genes (Figure 5C), it is possible that *de novo*suberin synthesis is induced in both incompatible and compatible interactions between plants and RKNs. This would be consistent with the fact that infection with RKN *M. incognita* induces the expression of *GPAT5*, a suberin biosynthetic gene, as well as the patchy suberin accumulation pattern within the cell layer surrounding infection sites in susceptible Arabidopsis plants (Holbein et al., 2019). Because the expression pattern of the suberin biosynthetic genes was similar to that of wound-inducible genes (Supplementary Figure 5), the induction of suberin biosynthesis could be triggered by wounding during nematode infection, but this need to be confirmed experimentally. In addition, it is important to clarify the biological relevance of ectopic suberin accumulation at the site of infection in the resistance response against PPNs in the future. We may be able to assess the individual contribution of ectopic suberin accumulation to the resistance response by selectively expressing a suberin-degrading enzyme CDEF1 (cuticle destructing factor 1) only at the site of infecton (Takahashi et al., 2010). However, transformation of *S. torvum* is technically challenging at the moment.

**FIGURE 8.**
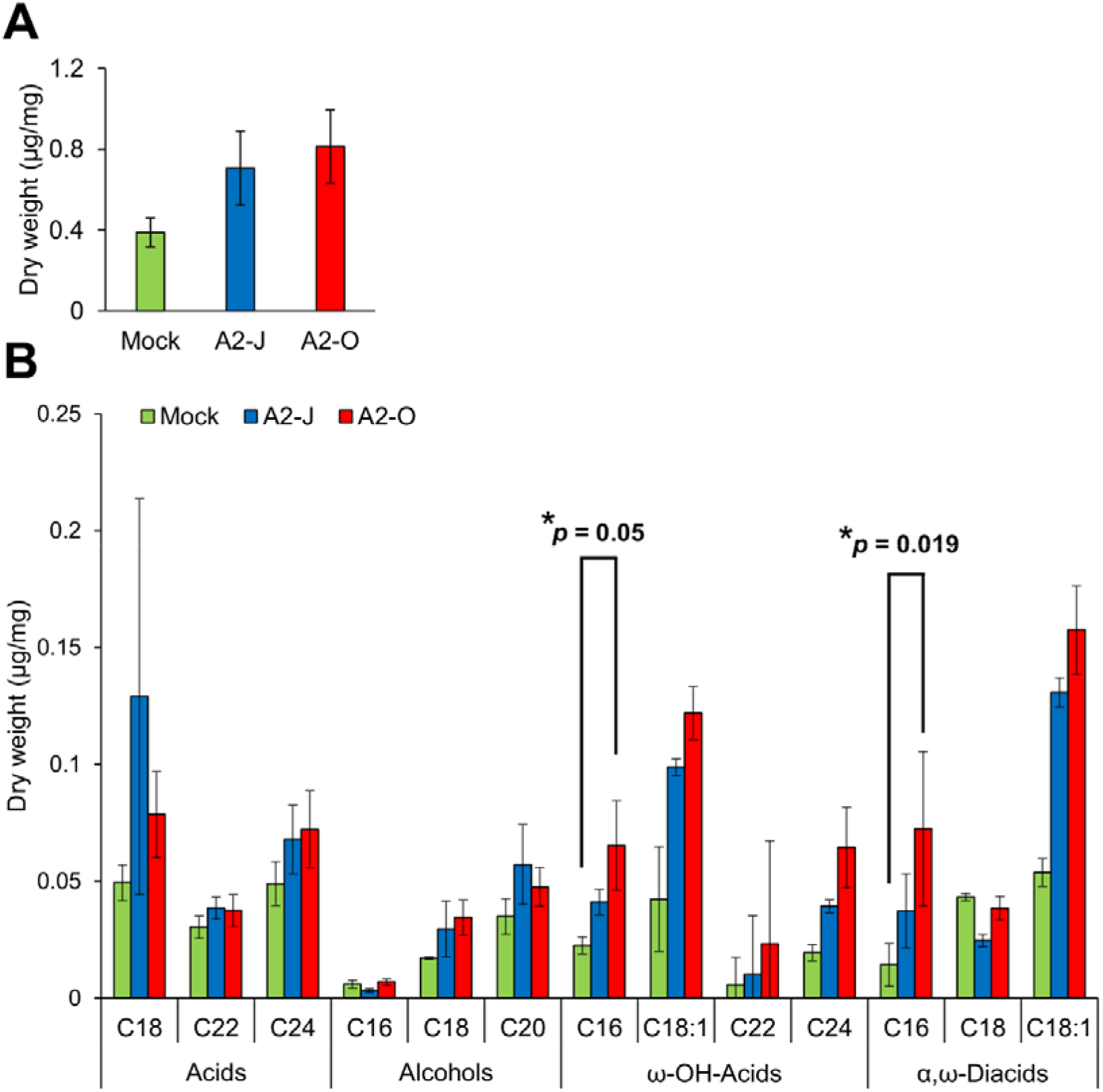
Infection with A2-O changed suberin monomer composition in the root tips. (**A**) Total aliphatic suberin and (**B**) suberin monomer composition in root tips treated with SDW (mock) or infected with A2-J or A2-O for four days. Gas chromatography-mass spectrometry (GC-MS) was performed to measure the content of aliphatic suberin monomers in root tips. Bars indicate means ± SE of four biological replicates. Asterisks indicate significant differences compared with mock treatment (Welch’s *t*-test with Bonferroni correction, * *p*≤ 0.05). There was no statistically significant difference at *p*< 0.05 in the total amount of aliphatic suberin between samples (Welch’s *t*-test with Bonferroni correction).

### A Model for Resistant and Susceptible Responses to RKN infection

*In this study, we were able to examine detail the transcriptional reprogramming of S. torvum in response to infection with virulent and avirulent RKNs at early infection stages by comparative transcriptome analyses (Figure 9). Shortly after infection with A2-O, S. torvum rapidly induces the expression of genes encoding fatty acid desaturases and sesquiterpene synthases, which may act as the first, or at least an early line of defense. Infection with A2-O also induces the expression of SA, JA, and ET marker genes, and defense-related genes (WRKY transcription factors*, chitinases, and β-1,3-glucanases), as well as the expression of lignin biosynthetic genes, leading to lignin accumulation in root tips. In contrast, infection with A2-J fails to alter gene expression at 1 DPI, but it does specifically induce the expression of genes encoding chalcone synthase and spermidine synthase, which might suppress lignin synthesis and the SA-mediated defense response (Hewezi et al., 2010). A2-J infection also induces the expression of genes related to cell wall modification and transmembrane transport, which may be important for the development and maturation of giant cells, and which in turn support nematode feeding, and are necessary to complete the parasitic life cycle. Interestingly, nematode invasion generally induces the expression of suberin synthetic genes, suggesting that the wounds caused by nematode entry and movement trigger suberin gene transcriptional activation. In the future, it will be important to independently assess the contribution of these genes to immunity and susceptibility against RKNs. It is also important to understand the molecular mechanisms and signal transduction pathways that decide the fate of infected plants and their nematode parasites, but will likely include whether or not plants have receptors that can recognize PPNs, and can transduce those signals to trigger defense responses, and/or whether or not PPNs produce virulence effectors that inhibit recognition, disrupt signal transduction, or suppress the defense response. As the next step, it is important to identify the specific effectors of A2-J and A2-O by comparative genomics, transcriptomics, and effectoromics, and to characterize their functions at the molecular level.

**FIGURE 9.**
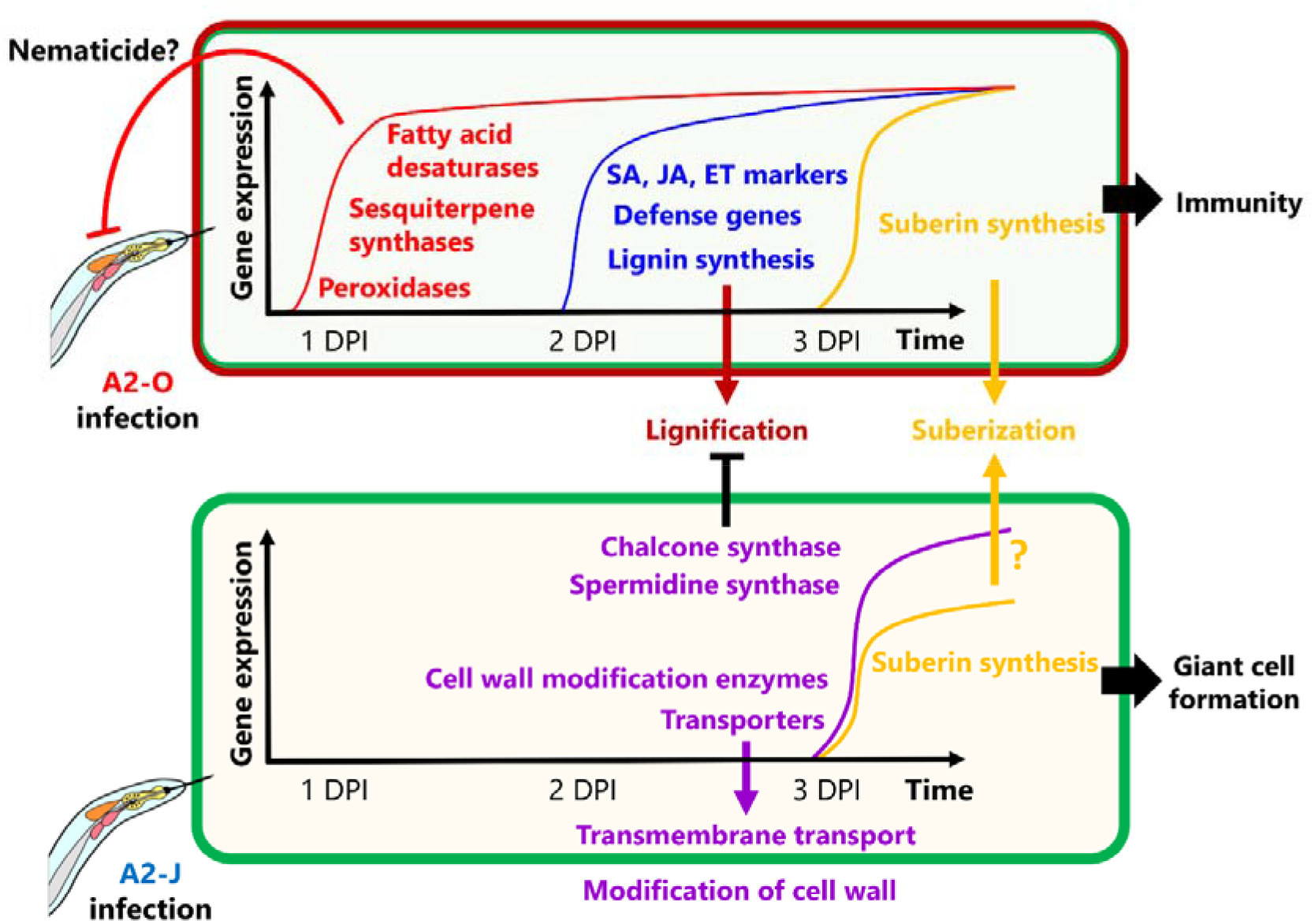
Schematic model of *S. torvum* responses to *M. arenaria* A2-O and A2-J. In the *S. torvum/M. arenaria* model, shortly after infection with avirulent RKN, plant genes encoding fatty acid desaturases and sesquiterpene synthases are induced to act as an early line of defense. Avirulent RKN infection also induces lignin accumulation in root tip, which may result in cell wall reinforcement that restricts RKN invasion and migration. Infection with virulent RKN results in a delayed gene expression response by specifically inducing chalcone synthase and spermidine synthase, which suppress multiple defense responses. Virulent RKN infection also results in induced expression of cell wall modification and transmembrane transport genes whose products contribute to giant cell formation for nematode feeding and completion of the life cycle.

## DATA AVAILABILITY STATEMENT

The transcriptome sequence data generated for this study are available in the National Center for Biotechnology Information, PRJNA681478. The *de novo* transcriptome assembly is available online at https://doi.org/10.6084/m9.figshare.14054354.

## AUTHOR CONTRIBUTIONS

Y.K. and K.Sh supervised the research. Y.K. designed the experiments. Y.K. and K.Sa carried out histopathological observation. Y.K. and T.U. propagate the nematodes and Y.K. collected samples for RNA-seq, lipid, and suberin analyses. N.M. generated RNA-seq libraries. T.S. carried out RNA-seq. Y.I. performed quality filtering of raw RNA-seq reads. K.Sa carried out bioinformatic analyses, including assembly, annotation, gene expression analyses, and GO enrichment tests. G.P. helped in bioinformatic analyses. T.B., K.Y., and S.Sh provided RKN genome assemblies. Y.K. and K.Sa performed lignin staining. Y.S.-S. and H.O. performed lipid quantification. J.H., R.B.F., S.Si, and F.M.W.G. carried out suberin monomer quantification. K.Sa, Y.K., and K.Sh wrote the manuscript. All authors read and approved the final manuscript.

## ACKNOWLEDGEMENTS

We thank all members of the Shirasu lab for intensive and helpful discussions. We thank Ms. Akiko Ueno, Ms. Naomi Watanabe, Ms. Mamiko Kouzai, Ms.Yoko Nagai, and Ms.Kanako Hori for their support of this project. Dr. Hideaki Iwahori at Ryukoku University for sharing *M arenaria* A2-O. We thank Mitsuyasu Hasebe, Shoko Ohi, Miwako Matsumoto (NIBB), and Tomoko Shibata for the PacBio sequencing and the Data Integration and Analysis Facility at NIBB for computer resources. We thank Dr. B. Favery for sharing the BABB cleaning protocols.

## Abbreviations

CN: Cyst nematode;
DPI: day-post inoculation;
RKN: Root-knot nematode;
PPN: plant-parasitic nematode

---

**SUPPLEMENTARY FIGURE 1.**
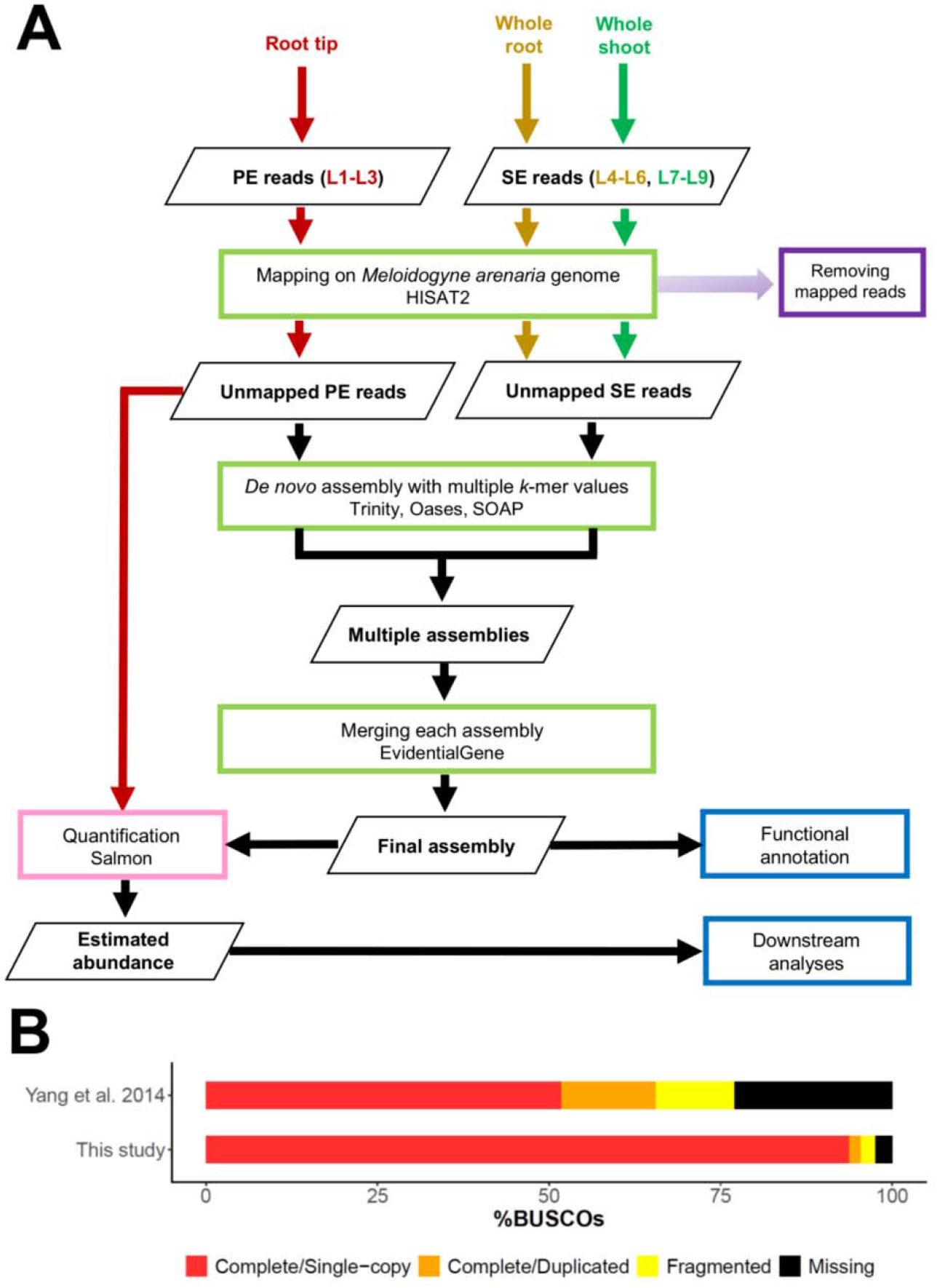
*In silico* analysis pipeline and completeness assessment of transcriptome assembly. (**A**) RNA-seq and bioinformatics pipeline. Paired-end (PE) reads for the root tip samples and single-end (SE) reads for the whole root and shoot samples were obtained from *S. torvum*treated with SDW (mock) or infected with A2-J or A2-O. PE and SE reads were mapped to the genome of *M. arenaria* A2-J or A2-O. Unmapped PE and SE reads were used for *de novo*transcriptome assembly. Unmapped PE reads were used for the quantification of transcript abundance. (**B**) BUSCO completeness comparison between this study and a previous study (Yang et al. 2014, BMC Genomics 15, 412) based on the Embryophyta odb9 dataset.

**SUPPLEMENTARY FIGURE 2.**
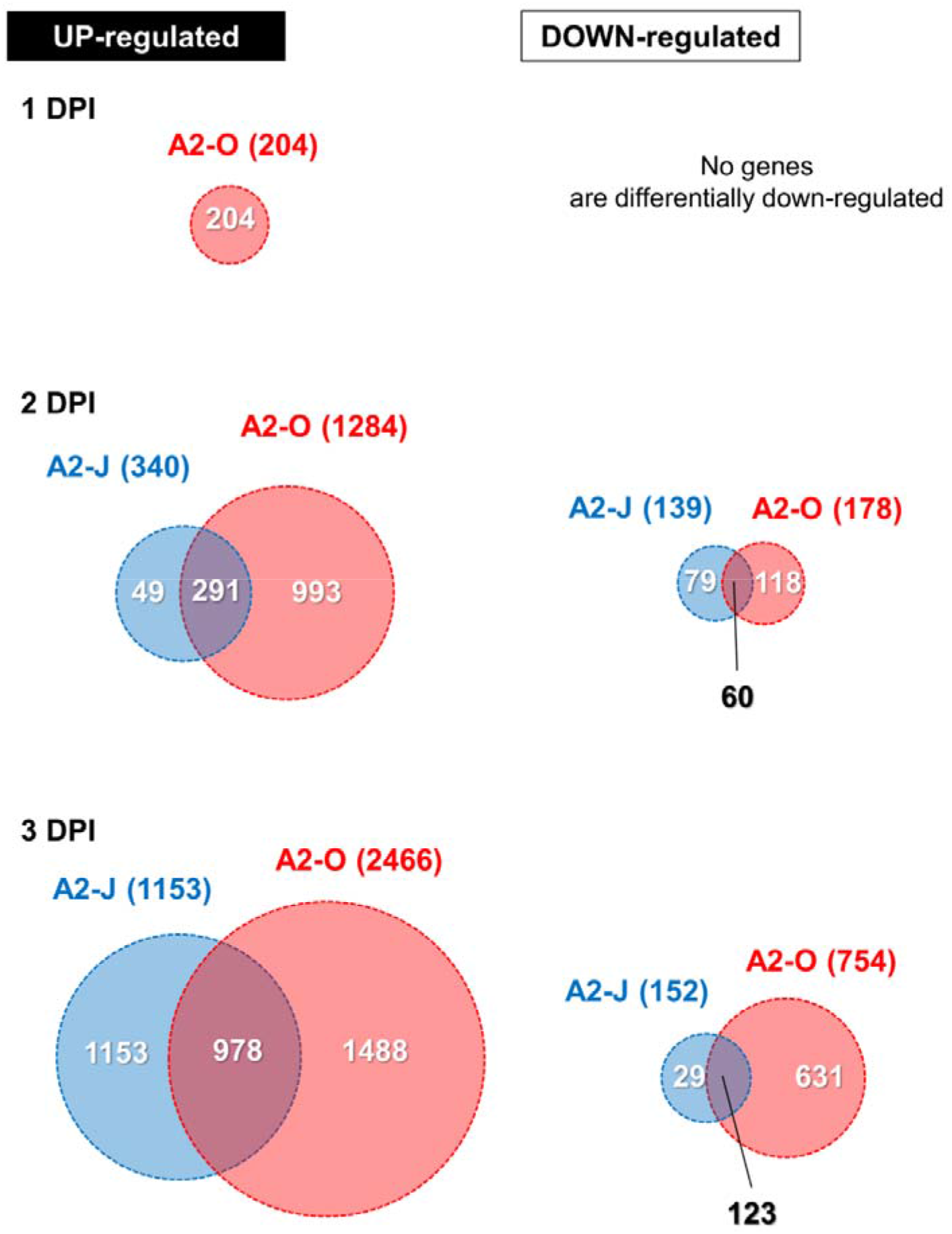
Differentially expressed genes in *S. torvum* infected with *M. arenaria* A2-J or A2-O. Venn diagrams showing the overlap of up-regulated genes (logFC ≥ 1, FDR ≤ 0.01) and down-regulated genes (logFC ≤ −1, FDR ≤ 0.01) at each time point after infection with A2-J and A2-O.

**SUPPLEMENTARY FIGURE 3.**
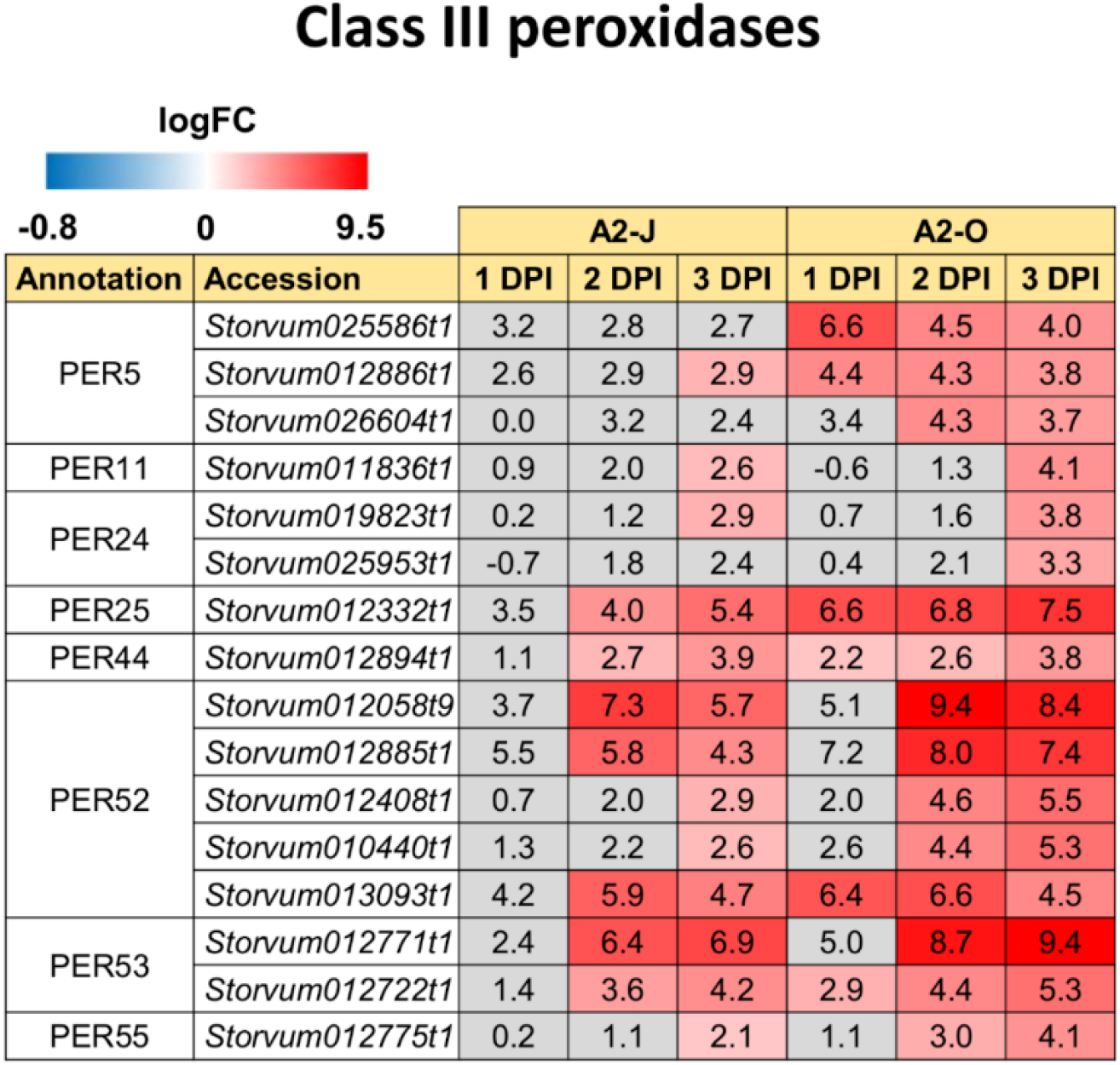
Infection with A2-O rapidly induced the expression of genes encoding class III peroxidase. Class III peroxidases in *S. toruvum* were searched based on Arabidopsis genome annotation (TAIR10). Highly up-regulated peroxidase genes (logFC ≥ 3, FDR ≤ 0.01) are listed. logFC values of the genes (compared to mock treatment) at 1, 2, and 3 DPI are shown. The heatmap uses grey to indicate values with no statistically significant differences at |logFC| ≥ 1 and FDR ≤ 0.01.

**SUPPLEMENTARY FIGURE 4.**
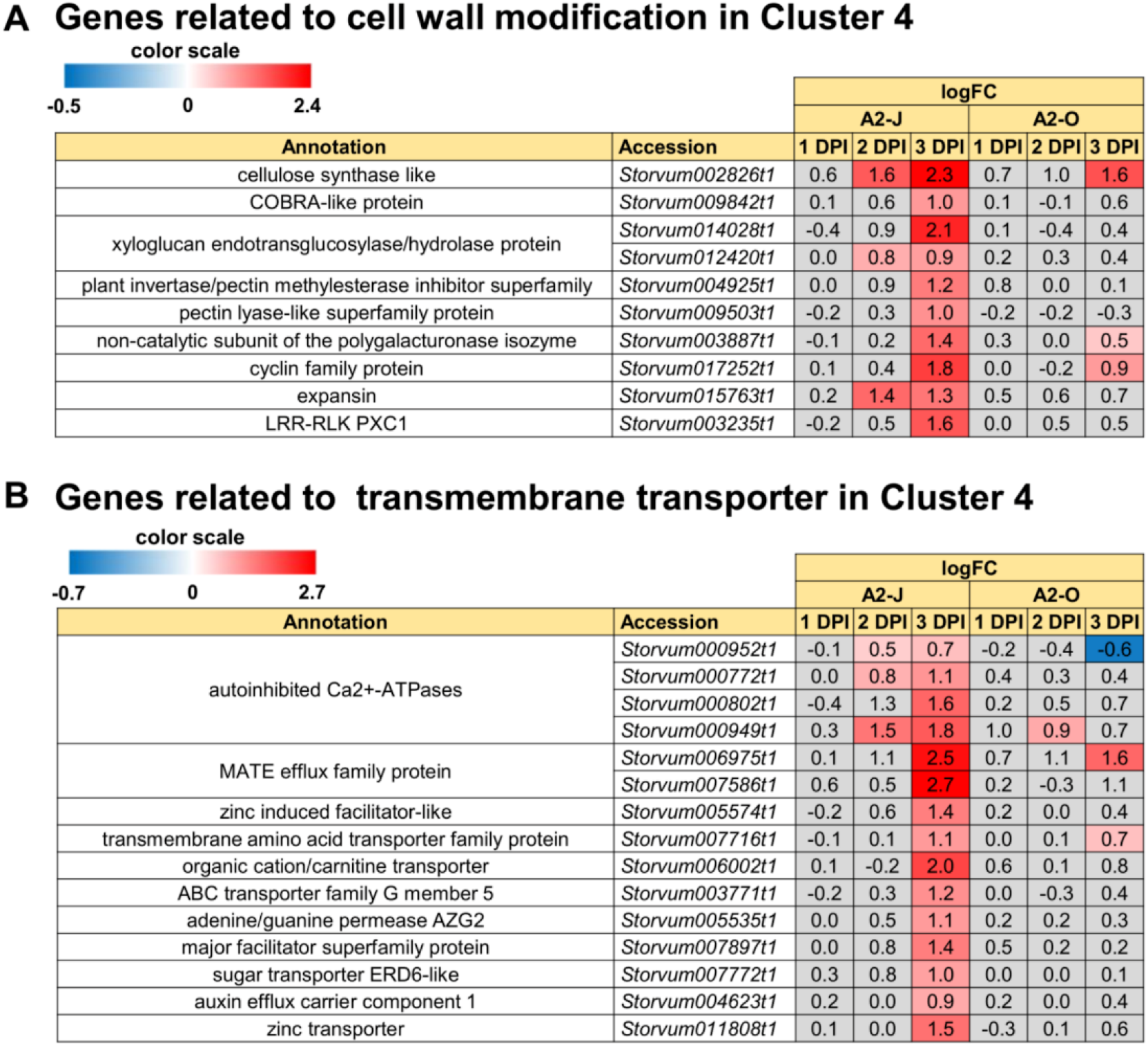
Infection with A2-J specifically induced the expression of genes related to cell wall modification and transmembrane transport. (**A**) The expression of genes related to cell wall modification in Cluster 4. Genes involved in cell wall modification in Cluster 4 were found in comparison with the gene lists of “cell wall modification (GO:0042545)”, “cell wall organization or biogenesis (GO:0071554)”, “pectin catabolic process (GO:0045490)”, and “polysaccharide metabolic process (GO:0005976)”. (**B**) The expression of genes related to transmembrane transport in Cluster 4. Genes involved in transmembrane transport in Cluster 4 were found by comparison with the gene lists of “transmembrane transport (GO:0055085)”. logFC values of the genes (compared to mock treatment) at 1, 2, and 3 DPI are shown. The heatmap uses grey to indicate values with no statistically significant differences at |logFC| ≥ 1 and FDR ≤ 0.01.

**SUPPLEMENTARY FIGURE 5.**
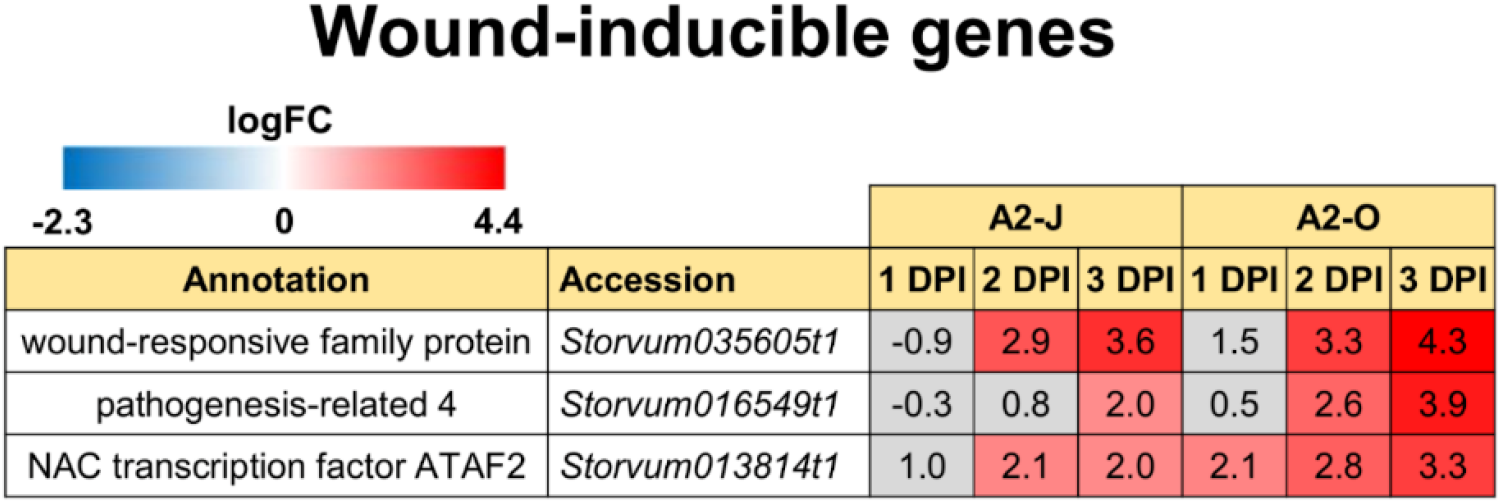
Both A2-J and A2-O induced the expression of wound-inducible genes. The expression of wound-inducible genes such as wound-responsive family protein, pathogenesis-related 4, and NAC transcription factor ATAF2 are shown. logFC values of the genes compared to mock treatment at 1, 2, and 3 DPI are shown. The heatmap uses grey to indicate values with no statistically significant differences at |logFC| ≥ 1 and FDR ≤ 0.01.

**SUPPLEMENTARY FIGURE 6.**
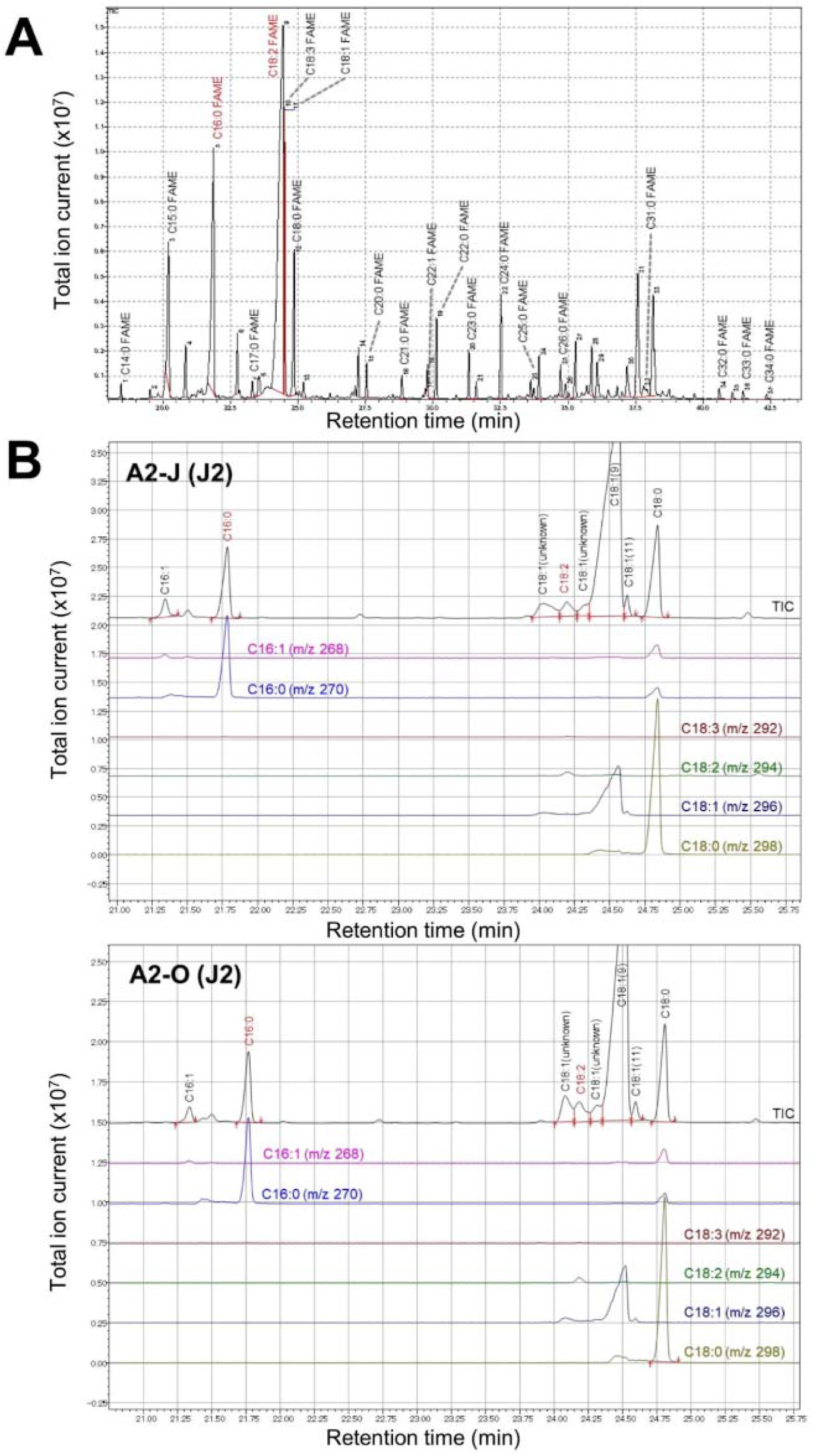
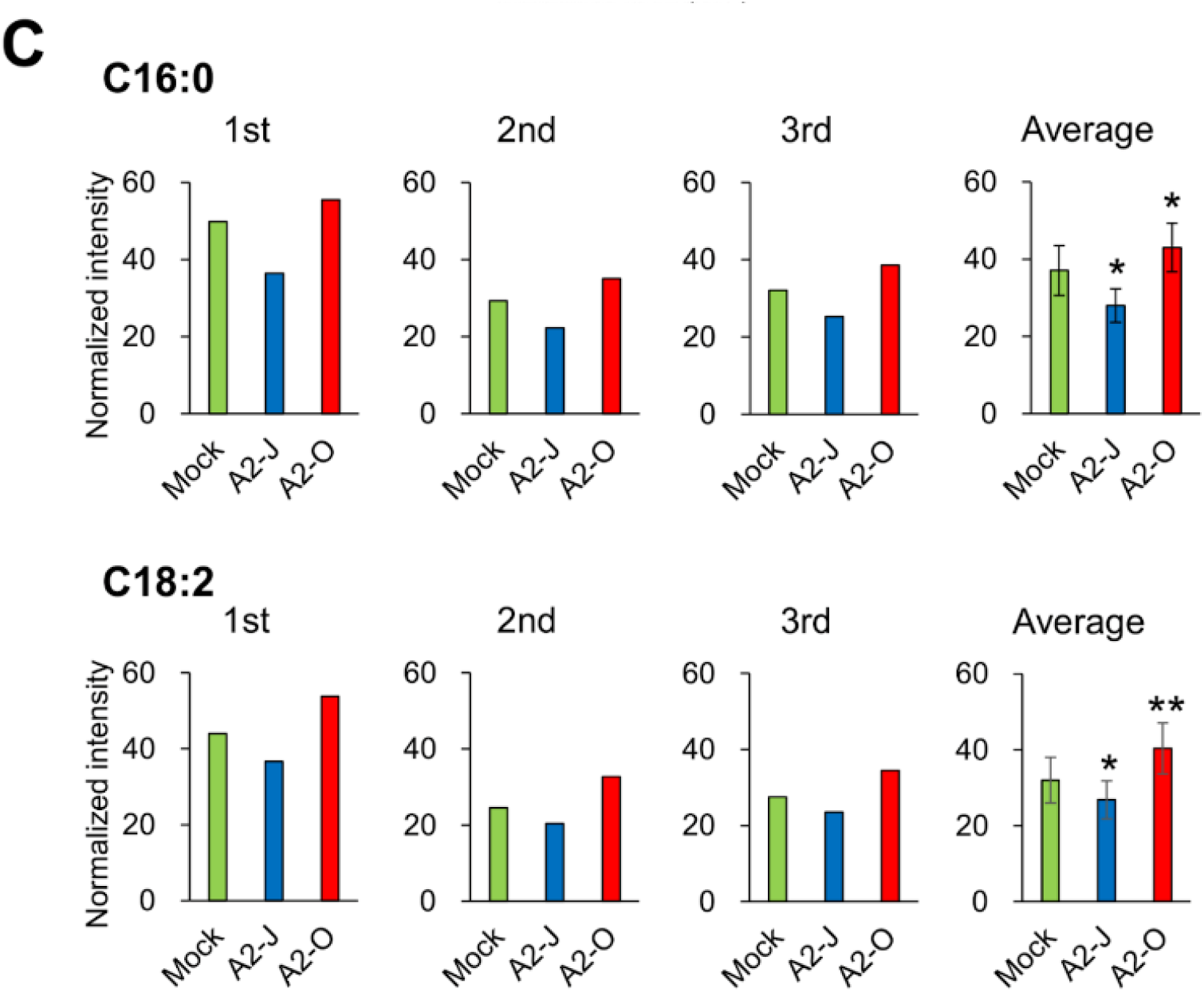
Infection with A2-O induced the accumulation of C18:2, and infection with A2-J reduced C16:0 and C18:2. One difficulty for measuring changes in the amount of each fatty acid in the root tip after infection with A2-J or A2-O is that it is not possible to distinguish an increase of particular fatty acids in root tips from contamination with the same fatty acids in nematodes. Thus, we looked for fatty acids that are abundant in root tips but are rare in nematodes. Total ion chromatograms (TIC) of root tips of *S. torvum* (**A**) and second stage juveniles (J2) of *M. arenaria* A2-J or A2-O (**B**) were obtained by gas chromatography-mass spectrometry (GC-MS). By comparing each peak, we found that C18:2 fatty acid methyl ester (FAME) was abundant in root tips, but was quite low in nematodes. C16:0 FAME was abundant in both root tips and nematodes. C16:0 and C18:2 in infected root tips (**C**). Data for each biological replicate and mean ± SE of three biological replicates are shown (one-way ANOVA, Dunnett post hoc test, * *p*≤ 0.05; \*\**p* ≤ 0.01). Infection with A2-O induced the accumulation of C18:2, infection with A2-J decreased C18:2 and C16:0 at 4 DPI. After infection with A2-O, C16:0 increased, but the higher amount of C16:0 may be due to the contamination of C16:0 originating in A2-O, but not due to active synthesis in *S. torvum*.

