## Supplementary material for "Transcriptomic analysis of resistant and susceptible responses in a new model root-knot nematode infection system using *Solanum torvum* and *Meloidogyne arenaria*": SUPPLYMENTARY MATERIALS AND METHODS

### Extraction and Preparation of Fatty Acid Methyl Esters (FAMEs)

J2s of RKN (*M. arenaria* A2-J and A2-O) and *S. torvum* root tips were treated with sterilized distilled water (mock) or infected with A2-J or A2-O (4 DPI), harvested, and flash frozen in liquid nitrogen. Total lipids were extracted as previously described (Bligh and Dyer, 1959). Lipids were dissolved in chloroform/methanol (2:1, v/v) and stored at -20 °C. FAMEs were obtained by incubating lipids for 1 h at 85 °C in the presence of 5 % (v/v) hydrogen chloride–methanol solution (Wako Pure Chemical Industries, Japan) (Iwai et al., 2014), then extracted with hexane and determined by gas chromatography-mass spectrometry (GS-MS).

### GC-MS Analysis

The qualitative composition of FAMEs was studied using a GS-MS (model GCMS-TQ8030; Shimadzu Corporation, Japan). High-grade pure helium was used as the carrier gas. The ionization voltage was 70 eV, and the ionization temperature was 200 °C. Mass spectra were scanned every 0.2 s. For the analysis of the FAMEs, a Rxi-5HT column (length, 15 m; internal diameter, 0.25 mm; film, 0.1 μm; Restek, PA, USA) was used. For each sample, 1 μL was injected onto the column into a helium gas flow held constant at 1.4 mL $min^{-1}$. The column temperature was elevated from 40 °C to 320 °C at a rate of 6 °C $min^{-1}$ and then maintained at 320 °C for 1.15 min.
